# Alpha and beta-diversities performance comparison between different normalization methods and centered log-ratio transformation in a microbiome public dataset

**DOI:** 10.1101/2022.11.07.512066

**Authors:** David Bars-Cortina

## Abstract

Microbiome data obtained after ribosomal RNA or shotgun sequencing represent a challenge for their ecological and statistical interpretation. Microbiome data is compositional data, with a very different sequencing depth between sequenced samples from the same experiment and harboring many zeros. To overcome this scenario, several normalizations and transformation methods have been developed to correct the microbiome data’s technical biases, statistically analyze these data more optimally, and obtain more confident biological conclusions. Most existing studies have compared the performance of different normalization methods mainly linked to microbial differential abundance analysis methods but without addressing the initial statistical task in microbiome data analysis: alpha and beta-diversities. Furthermore, most of the studies used simulated microbiome data. The present study attempted to fill this gap. A public whole shotgun metagenomic sequencing dataset from a USA cohort related to gastrointestinal diseases has been used. Moreover, the performance comparison of eleven normalization methods and the transformation method based on the centered log ratio (CLR) has been addressed. Two strategies were followed to attempt to evaluate the aptitude of the normalization methods between them: the centered residuals obtained for each normalization method and their coefficient of variation. Concerning alpha diversity, the Shannon-Weaver index has been used to compare its output to the normalization methods. Regarding beta-diversity (multivariate analysis), it has been explored three types of analysis: principal coordinate analysis (PCoA) as an exploratory method; distance-based redundancy analysis (db-RDA) as interpretative analysis; and sparse Partial Least Squares Discriminant Analysis (sPLS-DA) as machine learning discriminatory multivariate method. Moreover, other microbiome statistical approaches were compared along the normalization and transformation methods: permutational multivariate analysis of variance (PERMANOVA), analysis of similarities (ANOSIM), beta-dispersion and multi-level pattern analysis in order to associate specific species to each type of diagnosis group in the dataset used. The GMPR (geometric mean of pairwise ratios) normalization method presented the best results regarding the dispersion of the new matrix obtained after being scaled. For the case of *α* diversity, no differences were detected among the normalization methods compared. In terms of *β* diversity, the db-RDA and the sPLS-DA analysis have allowed us to detect the most meaningful differences between the normalization methods. The CLR transformation method was the most informative in biological terms, allowing us to make more predictions. Nonetheless, it is important to emphasize that the CLR method and the UQ normalization method have been the only ones that have allowed us to make predictions from the sPLS-DA analysis, so their use could be more encouraged.

## INTRODUCTION

The microbiome is a noun composed of micro and biome (both from Ancient Greece origin), meaning small and life, respectively. Even today, there are different microbiome definitions, depending on the scientific field of interest. However, based on the work of Berg et al. 2020 [1], a holistic definition of the term microbiome could be considered as follows: the microbiome is the sum of the microorganisms and their genomes in a particular ecological environment.

On the other hand, the microbiota concept integrates all the biological living forms that are part of the microbiome in a given ecological environment. In other words, if we are interested in the microbiome of the human gastrointestinal tract, we will talk about the concept of the human intestinal microbiota. On the other hand, if we are interested in the microbiota of a particular region of a National Park, we will use the term soil microbiota to refer to it.

The microbiome is made up of bacteria, fungi, algae and protozoa. Viruses, bacteriophages, and other mobile genetic elements, because they are not living beings [2], cannot be included in the definition. However, there is still controversy as to whether or not to include them [1].

To study the microbiome of the human gastrointestinal tract, non-invasive processes are usually recommended: stool collection (a representative sample of the intestinal microbiota) and saliva collection (a representative sample of the oral microbiota) [3]. From them, and without going into detail because it deviates from the objective of this work, they are subjected to a laboratory process of extracting the bacterial DNA they contain. Once extracted the genetic material, libraries are prepared to perform the sequencing, using two major technologies: 16S rRNA sequencing and shotgun metagenome sequencing [4–6]. In summary, and thanks to the cost reduction of shotgun technologies, the use of 16S sequencing is reducing because its main limitation is its taxonomic resolution: in the vast majority of cases, it can only be reached up to the taxonomic range of genus [7] and not to species. As a result of the sequencing process and its subsequent bioinformatic analysis (not described, out of scope), a matrix of counts is obtained, for example, at the species level for each sequenced sample. At this point, the microbiome dataset becomes compositional data and has been and continues to be a major headache for the applied statistics research field [8].

### Compositional Data. Theorical foundation

The compositional data is a matrix of non-negative numbers, with ***I*** rows and ***J*** columns, denoted by **X** (*I x J*). By convention, the rows ***I*** are the observation units (e.g. patients), *i*=1,2,…, *I*, while the columns (***J***) are the compositional parts (species in our example), *j*=1,2,…, *J*. Furthermore, by definition, the compositions in the rows of **X** are closed (sum up 1): 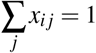, for all ***I***.

However, as a general definition, we can establish that a data set is compositional when the sum of the values for each sample are predefined [9]. The original values, whatever they were, are generally not of interest; instead, the relative values, collectively called composition, are relevant to understanding the structure of the data set. The components of a composition are called composition parts. If a subset of the parts is considered and the data is relativized relative to the new subtotals, this is called a sub composition [10].

The fact that the sum of the compositional data is constant makes it special. In any other more typical situation when the data has been collected on several variables (e.g., the amount of selenium, calcium, and bicarbonates in different commercial brands of mineral waters), there is absolutely no restriction on the value that each variable can have in each observation. In summary, each measurement collected is free to have a specific value in its particular measurement scale (unit of expression). By contrast, in compositional data, such as the count of microorganisms per patient, this freedom does not exist since they present the constant sum constraint. Generally, this constant sum is defined as 1 or 100%, although the original data is expressed, for example, in species counts (microbiome study) [11]. To show a basic example of compositional data, let us suppose that we ask four individuals to indicate how much time they dedicate to each activity (expressed in hours) on a specific day. In this case, the constant sum constraint will be 24 hours, the hours in a day (see Table 1).

**Table 1.** Example of compositional data on activities (in hours) in one day for four individuals. In this example, we have six variables, six components or six parts (sleep, meals,…) in the terminology used for compositional data.

| Individual | Sleeping | Meals | Work | Hobbies | Volunteering | Others | Sum |
| --- | --- | --- | --- | --- | --- | --- | --- |
| 1 | 8 | 1 | 8 | 3 | 1 | 3 | 24 |
| 2 | 9 | 1.2 | 9 | 0 | 0 | 4.8 | 24 |
| 3 | 7.5 | 1 | 5 | 6 | 2 | 2.5 | 24 |
| 4 | 8 | 1 | 4 | 4 | 4 | 3 | 24 |

Dividing a data set by its total to obtain the compositional values, which are proportions and sum up to 1, is called data closure or closure. Thus, having quantified the number of hours in the six-part composition of daily activities (see Table 1), the data would be closed (i.e., divided by 24 in this case) to obtain the values as proportions of the day.

Three principles define the analysis of compositional data, and they should be followed as closely as possible. These are: scale invariance, sub composition coherence, and permutation invariance [11].

The scale invariance principle states that compositional data only present relative (not absolute) information. Therefore, if we multiply the original data by any scalar factor ***C***, the compositional data remains the same after its closure.

The sub composition coherence principle means that results obtained for a subset of parts of a composition (known as a sub composition), should remain the same as in the composition.

Finally, the principle of permutation invariance means that the results do not depend on the order of the parts (variables) that appear in the composition. Of course, in a compositional data set, the parts are all ordered in the same way for each sample (individual), but the parts could be re-ordered without affecting the results.

The graphical representation of the compositional data helps its interpretation. The constant-sum constraint characteristic of compositional data causes the compositional data to have a special geometric representation of the compositions in a space known as the simplex. The simplest form of a simplex is a triangle (Figure 1), which contains three compositional parts (3 variables). A tetrahedron in three dimensions can represent 4-part compositions. Higher dimensional simplexes (with more than four compositions) are already challenging to represent.

**Figure 1.**
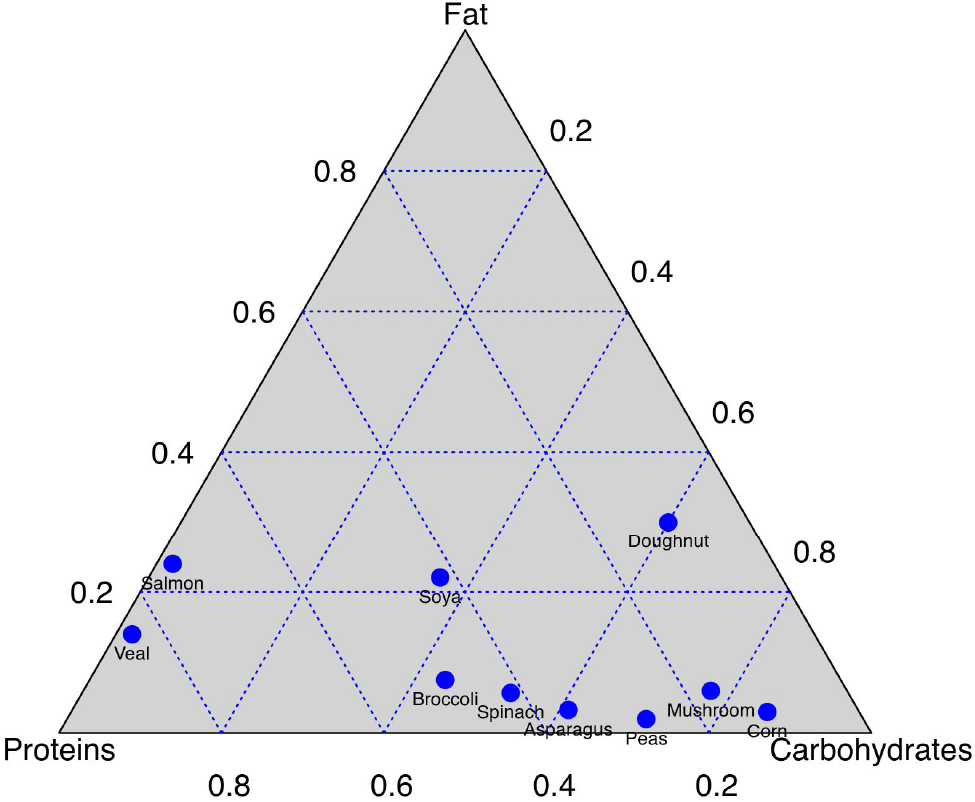
Compositon of three parts. Ternary graph. The variables are indicated at the vertices of the triangle. Most vegetables and doughnut, as opposed to veal and salmon, are high in carbohydrates. To discover it, draw the line parallel to the Fat-Carbohydrates axis that passes through the point of the food in question. The projection marks on the Proteins-Fat axis will indicate the percentage of Carbohydrates for that selected food. Figure adapted and recreated in R from Michael Greenacre’s book [11].

### Characteristics and challenges of the statistical analysis of the microbiome

The microbiome data, apart from being a microbiological species count matrix for each sequenced sample (compositional data), also has the following characteristics that make its subsequent statistical analysis difficult:

- Compositionality of the data
- Data sparsity
- High variability of the data
- Sequencing depth

Next, we will develop each of these last three characteristics since compositionality has already been explained in the previous section.

One of the characteristics of microbiome data, compared to other compositional data, is its big dispersion. The microbiome data contains many null values, meaning that species has not been detected for the considered sample x. This percentage of zeros in the microbiome data can be very high. Specifically, some human microbiome studies showed more than 80% zeros [12, 13]. This fact will make it challenging to find an adequate strategy for zero replacement [14] and is currently a very active field of research with the emergence of new strategies to combat the large data dispersion of the microbiome [15]. Microbiome data is sparse because several detected taxa are rare (uncommon) in the analyzed samples. Each sample has a unique microbiome composition, and only a few bacterial taxa will be shared among most of the analyzed samples. The rest will be rare taxa and only detected in small proportions [16, 17]. So far, only true zeros have been commented on (that particular taxon does not exist in the analyzed sample). However, among the zeros obtained in the data matrix, there also exists sampling zeros (null values due to sequencing depth inefficiency, discussed later) and technical zeros (null values due to the unwanted creation of experimental artifacts in pre-sequencing such as incomplete reverse transcription, polymerase chain reaction problems) [16, 18]. Because technical and sampling zeros cannot be distinguished from true zeros, all zeros are considered true [18].

Concerning the characteristic of the high variability of the data and its heterogeneity, it is an intrinsic property of the microbiome data. Microbiome data is a collection of counts from a long list of taxa that may have high and distinct levels of variability. For example, the set of abundant or rare taxa can vary considerably from sample to sample. The proportion of low or non-abundant taxa for most samples can be large (discussed in the previous paragraph). Like many other omics, microbiome data exhibit considerable natural heterogeneity or variability between samples. In addition to natural variability, there is also potential technical variation introduced by differences in sequencing depth and amplification biases [19, 20]. Regardless of the source, the total variability in microbiome data can be above and beyond what would normally be expected. The large variability coupled with excess zeros makes it difficult to identify true biological differences and can lead to biased estimation, and a high proportion of false positives [18].

Finally, the last fundamental characteristic that defines microbiome data is its sequencing depth, which has already been mentioned in previous paragraphs. This is a characteristic that is determined by the intrinsic limitations of the sequencing technology. Sequencing technology artificially limits the total number of counts observed per sample (also known as library size or sequencing depth). In other words, the counts of one taxon are directly affected by those of the others. Therefore, for a particular sample, an increase in abundance for one taxon means fewer available counts for all other taxa since the total number of counts cannot exceed the specified sequencing depth, which is limited by the sequencer capacity. The observed raw counts only reflect relative information, not the actual absolute abundances of the taxa in the samples (they are compositional data). In addition, due to the technical limitation of the sequencing depth that we have mentioned, we have that the total sum of the rows of the count table is not the same between the samples, which supposes an added difficulty in the statistical treatment of the microbiome or of other omics. In short, the sequencing depth is different for each sample. See the following Table 2 to understand better a real situation we can find in the microbiome analysis.

**Table 2.** Example of the compositional data in microbiome data. It can be seen that the sum (equivalent to the sequencing depth) is different for each sample, which adds an extra degree of difficulty to this type of compositional data.

| Individual | *E.coli* | *P.micra* | *B.oberum* | *M.smithii* | *H.massiliensis* | Sum |
| --- | --- | --- | --- | --- | --- | --- |
| 1 | 6500 | 1360 | 120000 | 0 | 0 | 127860 |
| 2 | 2000 | 540 | 11213 | 1345 | 50 | 15148 |
| 3 | 3500 | 5300 | 52360 | 6 | 1 | 61167 |
| 4 | 1237 | 150 | 3250 | 467 | 0 | 5104 |

Therefore, in summary, these four characteristics of the microbiome data described in the previous section make the first step before the statistical analysis of these data a challenge. This challenge process is called normalization, and it is an essential step as it will allow us to ensure or not the proper application of further statistical analysis [18]. Thus, the main goals of normalization techniques are: to remove any systematic technical biases, such as differences in sequencing depth or amplification biases that can negatively affect comparisons between samples; to take into account the large number of zeros that can be present in the data matrix; and try to make the observed counts as close as possible to the absolute counts (that is, transform the relative information of the compositional data into absolute terms).

In the present work, the main normalization techniques most used in microbiome data analysis will be presented. In addition, we will focus on alpha and beta-diversity since normalization methods in literature have been more focused to differential abundance analysis (see a highly recommended reading [18]), but not at the initial statistical analysis of how could affects the estimation of the diversities of the microbiome population studied. Finally, the comparison of the normalization methods has been carried out on a public microbiome data set and not on simulated data (a common strategy used in the bibliography consulted and referenced throughout the work).

## METHODS

### Bibliographic search

To gather information about microbiome data normalization methods, bibliographic references on compositional data have been used. A systematic search has been carried out until July 24, 2022, in PubMed database using the following keyword: “microbio* compositio*”, filtering for revisions and from 2018 (to get the most current view possible).

The review articles found with the filters mentioned above were critically analyzed if they dealt with normalization methods, and if so, it was checked if they presented references to other articles that detailed a normalization method accompanied by a mathematical development and/or statistical and (if possible) validated on some set of data (simulated or real). If not, the article was discarded.

### Microbiome data used

A search has been made in the 3 main open access databases of processed microbiome data (after bioinformatic analysis): microbiomeDB [21] (https://microbiomedb.org/mbio/app), MG-nify [22] (https://www.ebi.ac.uk/metagenomics/, and GMrepo [23] (https://gmrepo.humangut.info/home).

Specifically, a public study has been chosen with the use of shotgun technologies that allows for determining the taxonomic range of species (commented in the Introduction). Finally, the study by Franzoza et al. 2019 [24] has been selected, which is a cohort study with 56 healthy individuals, 88 individuals with Crohn’s disease, and 76 individuals with ulcerative colitis.

The processed data is available under the PRJNA400072 project at the following GMrepo link: https://gmrepo.humangut.info/data/project/PRJNA400072. The raw data was downloaded from the NCBI SRA webpage: https://www.ncbi.nlm.nih.gov/sra/?term=PRJNA400072.

### Normalization methods

Three large groups of normalization methods can be distinguished [18, 25]:

- Rarefaction
- Scaling methods
- Log-ratio transformations (CODA school philosophy, started by statistician John Aitchison, that 2022 marks the 40th anniversary of his first publication on compositional data assessment).

#### Rarefaction

The rarefaction method is the oldest normalization strategy, and it comes from the discipline of Ecology for the calculation of species richness [26]. Rarefaction is a method that adjusts for differences in library sizes between samples to aid comparisons of alpha diversity and beta diversity [27, 28]. Alpha diversity measures the diversity of species within a sample, while beta diversity accounts for differences in species composition between samples [29].

Rarefaction involves selecting a specific number of samples equal to or less than the number of samples in the smallest sample and then randomly discarding reads from the largest samples until the number of remaining samples equals this threshold. Based on these equal-sized subsamples, diversity metrics can be calculated that can contrast ecosystems “fairly”, regardless of differences in sample sizes [27]. Therefore, rarefaction solves the problem of the different sizes of the sequencing libraries between the samples that make up the data. It is important to emphasize that the sum of sequencing library sizes is related to the overall throughput of a particular sequencing run. Therefore samples sequenced on different sequencing machines or platforms will typically differ significantly in the library sizes. Also, and here is the kit for the matter, in a single batch of sequencing, you would expect to get approximately equal library sizes for all samples. However, in reality, after sequencing, each sample is associated with a very different number of reads. The different numbers of reads for each sample reflect the differential efficiency of the sequencing process between samples (for example, uncertainties in library quantitation and/or variation in loading concentrations or volumes) rather than the biological variation of interest[28]. Therefore, and in microbiome data, we will encounter the problem of different library sizes between samples.

To facilitate its understanding, the rarefaction process is explained below again but in a more pleasant way, based on Hong et al. 2022 [28]:

Let ***L***✽ be the (arbitrarily) chosen sequencing library size:

1. Specify a library size (*L*^✽^ ≤ *max_i_*(*L_i_*)) where *i* indexes all samples;
2. Discard all those samples with library size *L_i_* < (*L*✻); and
3. For the samples left over from the previous step (*L_i_* ≥ (*L*^✽^)), separately for each sample, randomly subsamples its reads without replacement to ***L***^✽^.

The selected sequencing library size (***L***^✽^) for the set of samples considered is often chosen to be the smallest library size observed, assuming that all samples in question have been correctly sequenced (for example, ignoring/eliminating those samples that have very small library values compared to the rest of the samples, due to errors during sequencing discussed above) [28]. However, the larger (***L***^✽^) is, the amount of artificial variation introduced in diversity analyses is minimized but may require the omission of samples with small library sizes [30].

The classic R packages for microbiome analysis (phyloseq and vegan) incorporate functions to calculate rarefaction but only by performing a single iteration. Recently a new R package (myrlin) defaults to 1000 iterations [30]. In addition, this year a rarefaction index has been developed that, the closer its value is to 1, the rarefaction will not imply any distortion of the results [28], since the rarefaction has been criticized [28, 31] but is still used today as it is a method that works very well in the ecology and microbiology disciplines [28, 32].

#### Scaling methods

The following scaling methods will be described in this section: TSS, CSS, TMM, DESEq2, ELib-TMM, ELib-UQ, UQ, GMPR, Wrench, and ANCOM-BC. Scale normalization methods attempt to correct observed counts for systematic bias using a scale factor that is often sample-specific.

The scaling factor can be defined through the following Equation 1 where the basic idea is to divide the observed counts in the species table by a “scaling factor” (or “normalization factor”) to remove those biases due to sequencing depth difference [33].

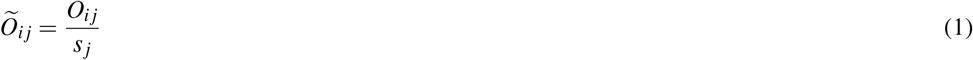

where 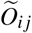 is the normalized observed abundance for taxon *i* for each sample *j; O_ij_* is the observed abundance of the i-jth taxon (species) in the j-th sample; and *s_j_* is the scaling/normalization factor for sample *j*.

Since a large part of the technical variability comes from the differences in the total reads per sample (sequencing library size), some commonly used normalizations such as Total Sum Scaling (TSS), Trimmed Mean of M-values (TMM), and Upper Quartile (UQ) attempt to correct for observed counts and compensate for differences in sequencing depths. Other methods, such as Wrench and the ANCOM-BC [33] attempt to provide additional scaling for data compositionality and sparsity [18].

The simplest and most direct scaling normalization method that corrects for differences in sequencing depth is the TSS method. TSS normalization scales individual read counts by the total number of reads, thus transforming the observed abundances into relative abundances. However, the relative abundances remain compositional since the sum total of abundances for a sample is set to 1. The TSS method scaling factor formula is presented in Eq. 2.

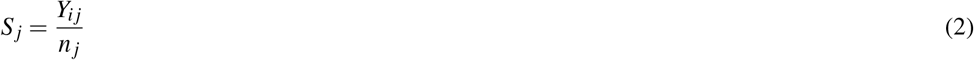

where *i* = 1*p* is the index of the taxon (eg species), *j* = *N* is the index of the samples (individuals); *Y_ij_* the untransformed counts of the i-th taxon in the j-th sample; *S_j_* is the scaling factor for the j-th sample; and *n_j_* the size of the sample library *j*.

The CSS method (Eq. 3) uses robust statistics to provide an alternative to TSS that is less influenced by preferentially sampled taxa. CSS is defined as the cumulative sum of observed counts up to a threshold that is determined using a heuristic that minimizes the influence of preferentially sampled taxa. Thus, CSS attempts to scale each sample using only the relatively invariant part of the count distribution [18]. However, neither TSS nor CSS do not take into account the compositionality of the data or the dispersion of the data.

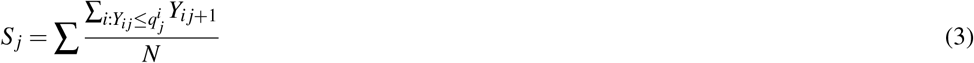

where *i* = 1,…,*p* is the index of the taxon (for example species), *j* = 1,…,*N* is the index of the samples (individuals); *Y_ij_* the untransformed counts of the i-th taxon in the j-th sample; *S_j_* is the scaling factor for the j-th sample; and 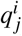 is the i-th quantile of the sample *j*.

The TMM method (Eq. 4), for each sample, chooses a reference that will be the weighted trimmed mean of the logarithmic abundance indices after exclusion of the most abundant taxa that have the highest values of log ratio, and uses this scaling factor to normalize the size of the corresponding library. Like the DESeq2 method, TMM assumes that most taxa are not differentially abundant.

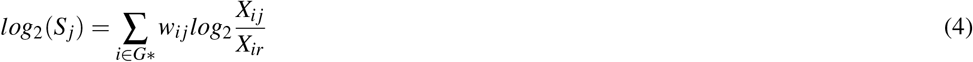

where *i* = 1,…, *p* is the index of the taxon (eg species), *j* = 1,…, *N* is the index of the samples (individuals); *X_ij_* represents the relative abundance of taxon *i* and displays *j; S_j_* is the scaling factor for the j-th sample; *G*^✽^ represents the trimmed set of taxa by *j; w_ij_* represents the specific weight for each method; and *X_ir_* is the reference sample for taxon *i*.

The DESeq2 method (Differential gene expression analysis based on the negative binomial distribution) (Eq. 5) chooses as reference for each taxon the geometric mean of the abundances in all the samples. The DESEq scaling factor of the observed abundances for each sample is calculated as the median of all ratios between the sample and reference counts.

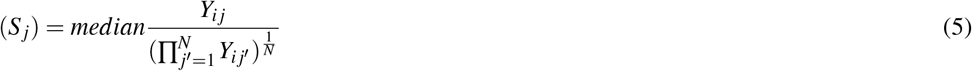

where *i* = 1,…, *p* is the index of the taxon (eg species), *j* = 1,…, *N* is the index of the samples (individuals); *X_ij_* represents the relative abundance of taxon *i* and displays *j*; and *S_j_* is the scaling factor for the jth sample.

The ELib-TMM method is a modified version of the TMM method that takes into account the corresponding library size for each sample (Eq. 6).

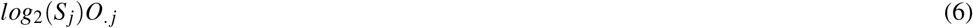

where *log*_2_(*S_j_*) is the result of TMM (Eq. 4) and 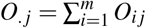 where *O_ij_* is the size of the library for the i-th taxon for the j-th sample; and *m* the total number of taxa.

The UQ method (Eq. 7) observations of each taxon are divided by the upper quartile of the (non-zero) counts associated with each sample and multiplied by the mean upper quartile of all the dataset samples [34].

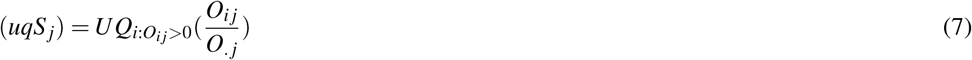

where *uqS_j_* is the scaling factor for the j-th sample, *UQ* is the upper quartile, and 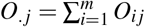 in which *O_ij_* is the size of the library for the i-th taxon for the j-th sample; and *m* the total number of taxa.

The ELib-UQ (Effective library size using UQ) method is a modified version of the UQ method that takes into account the corresponding library size (Eq. 8).

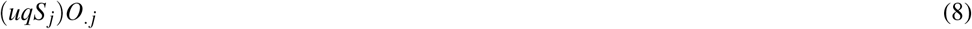

where (*uqS_j_*) is the result of UQ (Eq. 7) and 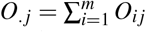 where *O_ij_* is the size of the library for the i-th taxon for the j-th sample; and *m* the total number of taxa.

The GMPR (geometric mean of pairwise ratios) method (Eq. 9) is a compositional normalization method specifically designed to deal with sparse data (eg, microbiome data) and takes into account the sizes of their libraries. It is a method that represents an extension to the DESEq2 method by reversing the steps of DESEq2 to deal with data sparsity *(sparsity).* First, the median of all pairwise proportions of counts ≠ 0 from two samples is computed. The scale factor for a sample is then calculated by combining the pairwise results for the sample to obtain the geometric mean of the median values for that sample and all other samples.

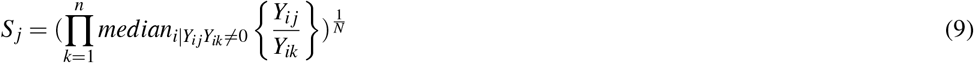

where *i* = 1,…, *p* is the index of the taxon (eg species), *j* = 1,…, *N* is the index of the samples (individuals); *X_ij_* represents the relative abundance of taxon *i* and displays *j; Y_ij_* the untransformed counts of the i-th taxon in the j-th sample; and *S_j_* is the scaling factor for the jth sample.

The Wrench method (Eq. 10) is considered a generalization of TMM for zero-inflated data *(zero-inflated data)* that reduces the estimated biases that occur with normalization methods that ignore the zeros (like the TMM). The Wrench method attempts to eliminate library size biases and allows absolute (not relative) observations to be obtained after normalization. To achieve this, it estimates a “compositional correction factor” which is the value that estimates the systematic bias in a group.

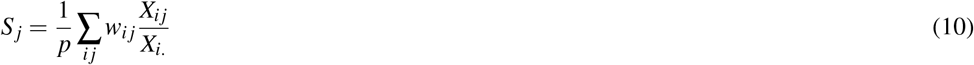

where *i* = 1,…, *p* is the index of the taxon (eg species), *j* = 1,…, *N* is the index of the samples (individuals); *X_ij_* represents the relative abundance of taxon *i* and displays *j; w_ij_* represents the specific weight for each method; and *S_j_* is the scaling factor for the jth sample.

The ANCOM-BC method (Analysis of Compositions of Microbiomes with Bias Correction) (Eq. 11) allows Wrench to infer absolute abundance from relative abundance. The authors of this method incorporate the term sampling fraction *(sampling fraction)* as the ratio of the expected observed abundance of the taxon in a random sample. See Figure 1 of the article by Huang Lin et al. 2020 [33] to delve into this interesting concept.

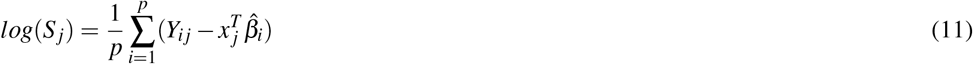

where *i =* 1,…, *p* is the index of the taxon (eg species), *j* = 1,…, *N* is the index of the samples (individuals); *Y_ij_* the untransformed counts of the i-th taxon in the j-th sample; 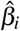 represents the estimate obtained in ANCOM-BC (sample fraction); and *S_j_* is the scaling factor for the jth sample.

#### Log-ratio transformations

Transformations are not proper normalization methods like the scaling and rarefaction methods discussed above. In the transformations, the counts are transformed based on a reference to perform statistical inferences based on the chosen reference [18, 25].

Within the wide range of data transformations that we could think of, the analysis methods to analyze compositional data (CODA) (method introduced by Join Aitchison [35]) use the log-ratio transformation. The most well-known and used log-ratio transformation in microbiota (and which we will focus on in this work) is the centered log-ratio (CLR) followed by the additive log-ratio (alr) and the inter-quartile log-ratio (iqlr) [**Street_2019**, 11, 18, 25].

The CLR transformation uses as a reference the geometric mean of the vector of each sample (Eq. 12).

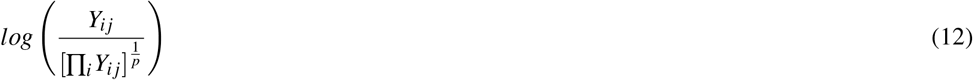

where *i* = 1,…, *p* is the index of the taxon (eg species), *j* = 1,…, *N* is the index of the samples (individuals); and *Y_ij_* the untransformed counts of the i-th taxon in the j-th sample.

Depending on the calculation of logarithms, CODA methods cannot compute zeros, which are very common in omics analyses, such as microbiota. To overcome this scenario, before calculating the CLR, a pseudo value must be added to all the zeros or (more recommended) an imputation of the zeros should be made based on a Bayesian multiplicative replacement strategy [11, 25].

### Statistical analysis

#### Statistical software

All data have been analyzed with the statistical software R version 4.1.2 [36] with its interface RStudio 2021.09.01 [37]. The SessionInfo() together with all the R script codes are available from Zenodo repository (https://doi.org/10.5281/zenodo.7134538).

#### Comparison between normalization methods and CLR transformation

For the evaluation of the different of normalization methods and CLR transformation, the following R libraries have been used: ANCOM-BC (ANCOM-BC package) [38]; CSS (metagenomeSeq package) [39]; DESeq2 (DESeq2 package) [40]; TMM, UQ, ELib-TMM and ELib-UQ (edgeR package) [41]; GMPR (GMPR package) [42]; rarefaction (phyloseq and mirlyn packages) [30, 43]; TSS (base package) [36]; Wrench (Wrench package) [44]; and centered log-ratio (easyCODA package) [11].

The 11 normalization methods have been compared with each other and to the original non-normalized data. As discussed and discussed below, the CLR method is not a normalization method [29][29] and its suitability for use in microbiome data has been evaluated in another way described in the next subsection. Two approaches have been made to compare the normalization methods between them: (i) comparing the centered residuals between normalized methods from the code available from [33, 38], and (ii) the comparison of the coefficient of variation (expressed in %) between the normalization methods obtained through the rowMeans functions () from the base package [36] and rowSds() from the matrixStats package [45] (see R code for further details).

#### Microbiome statistical analysis

To compare the results obtained using the different normalization and transformation methods for a public data set, alpha diversity (diversity of species in a specific sample analyzed) [46] has been evaluated using the Shannon-Weaver (Eq. 13) that takes into account both the abundance and the uniformity of the species [47] and is highly popular in microbiota studies [48]. For this, the R package used has been that of phyloseq [43] and ggplot2 [49].

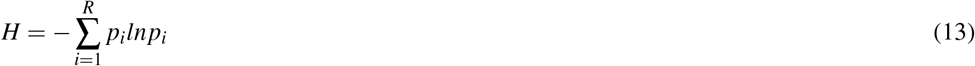

where *p_i_* is the proportion of individuals belonging to species i.

On the other hand, their possible differences in the beta-diversity of the studied community (difference in species between samples) [46] have been evaluated between the different methods using the Bray-Curtis distance (Eq. 14). Bray-Curtis distance is widely used in the fields of ecology, and microbiome [32, 50].

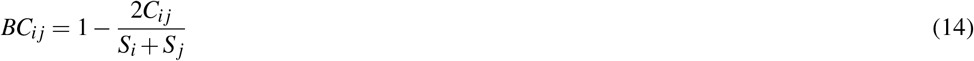

where given a sample *i* and a sample *j, C_ij_* is the sum of the lower values for those species common between the two compared samples. *S_i_* and *S_j_* are the total number of species for each sample.

For the statistical analysis of beta-diversity, the multivariate analysis is used [51]. Within the multivariate analysis, we distinguish three large groups: exploratory, interpretative, and discriminatory methods [52]. Exploratory methods are used to evaluate the relationship between objects based on the values of variables measured in that objects. Those similar objects will be distributed close to each other, while dissimilar objects will be distributed at non-close points on the graph. Interpretative methods are “constrained techniques”, which in addition to the main set of measured variables, also use another set of additional explanatory variables (for example, known environmental gradients between objects). This constrained ordination analysis aims to find axes in the space of the multidimensional data set that maximizes the association between the explanatory variable(s) and the measured variables (response variables). Therefore, the ordination axes are constrained to be functions of the explanatory variables. The coefficients for each explanatory variable used to calculate each ordination axis indicate the contribution of that variable to the dispersion of the observed object along that axis. Finally, the discriminatory methods aim to define discriminant functions (synthetic variables) that maximize the separation of the objects between the different classes. The discriminant function(s) are restricted to a specific combination of explanatory variables. Variable coefficients (also known as weight or *loadings*) are used to calculate each discriminant function, and they indicate the relative contribution of each explanatory variable to the observed separation of the object along each discriminant function [52].

It exists a wide range of multivariate analyzes applicable to ecology and to our particular case of the gastrointestinal microbiota. For pedagogical reasons and the length of the present suty, one type of technique has been selected from each type of multivariate analysis. Principal Coordinate Analysis (PCoA) has been selected as an exploratory method, which is an extension to Principal Component Analysis (PCA), but while PCA organizes the objects by an analysis of the eigenvalues of the correlation matrix, PCoA is can be applied to any distance measure, including the Bray-Curtis distance that we have used in this work. The PCoAs have been carried out through the phyloseq package [43].

As an interpretative method, the distance-based redundancy analysis (db-RDA) method has been chosen since it allows using the Bray-Curtis distance [53]. To analyze the db-RDA, the vegan [54] package has been used.

Finally, the sPLS-DA (sparse Partial Least Squares Discriminant Analysis) technique has been used as a discriminative method, which represents an improvement over PLS-DA because it allows for a selection of variables, discarding those variables that are not informative [55]. For the sPLS-DAs, the mixOmics package [56] has been used.

For the analysis of db-RDA and sPLS-DA in the set of additional explanatory variables that contained missing data, an imputation based on the random forest machine learning technique was performed through the package missForest [57]. In addition, to represent it correctly, all the categorical variables of the matrix of additional explanatory variables have been converted to dummy variables (with a value of 0 or 1) through the use of the fastDummies [58] package. Also, due to the overlapping of the species in the resulting graphs, and to achieve a more pleasant reading of them, a graph has been made with a subset of 47 species selected by the alternative method to SIMPER detailed later.

Regarding the case of the CLR transformation and according to the consulted bibliography [11, 29, 59], the beta diversity cannot be evaluated with the Bray-Curtis distance but Euclidean, then the PCoA transforms into a PCA [29]. To do this, the pco() function of the ecodist package [60] is used, having performed previously the imputation of zeros with the cmultRepl() function of the zCompositions package [61]. The other two types of multivariate analysis could be compared with the other normalization methods using the same analysis strategy. However, alpha diversity has not been studied with the CLR transformation because alpha diversity formulas only support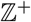 [43]. The R scripts used are found in Zenodo repository.

Finally, the differences between diagnoses (control, ulcerative colitis, and Chron’s disease) in the composition of the intestinal microbiota have been evaluated in a complementary way with the following approaches: (i) PERMANOVA through the adonis() function [54], (ii) analysis of similarities (ANOSIM) [54], (iii) the beta dispersion [54], and (iv) an alternative to SIMPER from the vegan package [54] (multipatt) of the indicspecies package [62] that has allowed to associate specific species to groups of specific diagnosis, unlike SIMPER that only allowed comparisons between groups of diagnoses. The multipatt function is a multilevel pattern analysis that calculates an indicator value for each species in association with the input groups and then finds the group with the highest association with each species. The statistical significance of the associations is then tested using the indicator value as the test statistic in a permutation test. The PERMANOVA posthoc test has been performed through the pairwise.perm.manova() function of the RVAideMemoire package [63].

## RESULTS

### Brief description of the analyzed data

As indicated in the Materials section, they have used public microbiome analysis data [24]. A brief summary of the general descriptive statistics of these data is indicated in the following Table 3. Of a total of 220 patients, 56 did not present a pathology associated with the digestive system while 88 presented Chron’s disease and 76 ulcerative colitis. The overall mean age was 43 years and information was collected on the content of calprotectin in feces (numerical variable) and the consumption or not of antibiotics, immunosuppressants, mesalamine and steroids (categorical variables).

**Table 3.** Statistical summary of the analyzed data. CD: Crohn’s disease. CU: ulcerative colitis. NR: not collected/answered.

| Individuals | Diagnostic | Age | Calprotectin | Antibiotic | Immunosupressants | Mesalamine | Steroids |
| --- | --- | --- | --- | --- | --- | --- | --- |
| 220 | Control: 56 | Min: 19 | Min:0.67 | No:199 | No:131 | No:133 | No:157 |
|  | CD: 88 | Mean: 43 | Mean: 146 | Si:18 | Si:67 | Si:63 | Si:39 |
|  | UC: 76 | Max: 82 | Max:2440 | NR:3 | NR:22 | NR:24 | NR:24 |

### Comparison between normalization methods

As has been commented in the Material and methods section for the comparison between the normalization methods, a public data set it has been used (they are not simulated data), and two strategies have been followed. The first strategy has been to compare the centered residuals through a box plot between the true sample fraction and its estimate for each sample. See Figure 2. In this first strategy, only the scaling-type normalization methods have been compared. From the graph, we can indicate that the most desired box plot output will be the one with a lower height (less variability) and no (or few) peripheral points. Consequently, the most recommended scaling methods for our analyzed public data would be ANCOM-BC, TSS, ELib-TMM, GMPR and DESeq. ELib-UQ presents the smallest height of its box but presents a high variability (3.7) and many “peripheral” points. It is also noteworthy that there has been no scaling normalization for the residuals to appear grouped according to the diagnosis.

**Figure 2.**
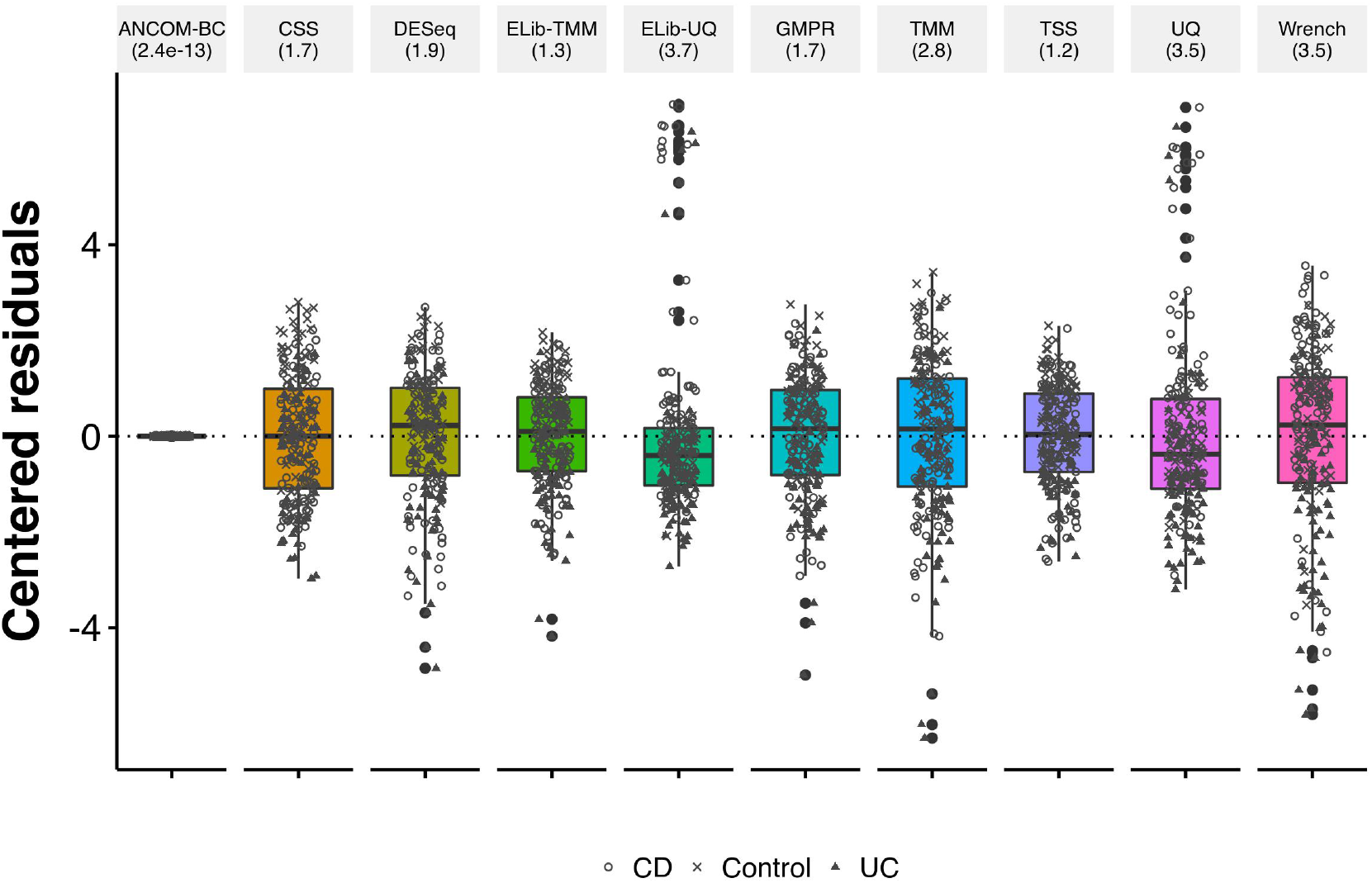
Box plot of the residuals between the true sample fraction and its estimate for each sample. The lower and upper hinges correspond to the first and third quartiles (25th and 75th percentiles, respectively). The median is represented by the solid black line inside the box. The upper whisker extends from the hinge to the largest value no more than 1.5 times the interquartile range (distance between the first and third quartiles). The equivalent in the lower whisker. Data beyond the whiskers is called “outer” points. The value in parentheses associated with each normalization method is the variance.

In the second strategy (see Figure 3), all the normalization methods have been taken into account, and the original data without normalizing. We can see that three methods (rarefaction, TSS (equivalent to calculating relative abundances), and UQ) have presented identical results. However, the methods that have presented a better aptitude (based on the criteria discussed in the previous figure) compared to the original data (NOT_NORM) have been Wrench, TMM, and GMPR.

**Figure 3.**
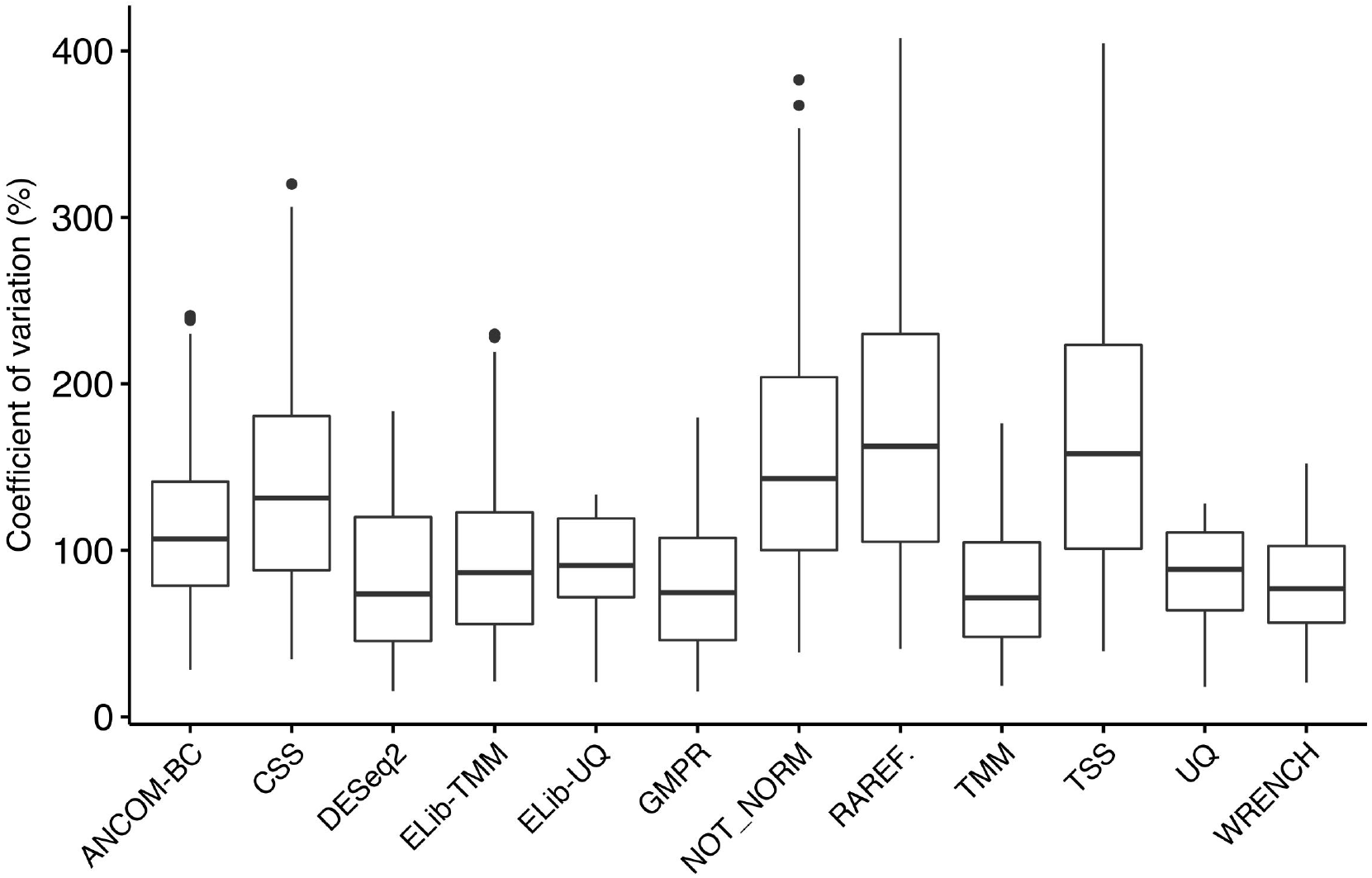
Box plot of the coefficients of variation in the different normalization methods studied. The lower and upper hinges correspond to the first and third quartiles (25th and 75th percentiles, respectively). The median is represented by the solid black line inside the box. Data beyond the whiskers are possible *outliers.*

### α diversity

The Shannon-Weaver index has been selected to compare the *α* diversity between the different normalization methods (Figure 4). The TSS, ELib-TMM, and ELib-UQ normalization methods do not appear for the reason mentioned in Materials and Methods. Although some small differences were detected in the p-values (Wilcoxon test) between the control group and CD (Chron’s disease), all the normalization methods converge to the same results, and there are no differences in the Shannon index according to the normalization method employed.

**Figure 4.**
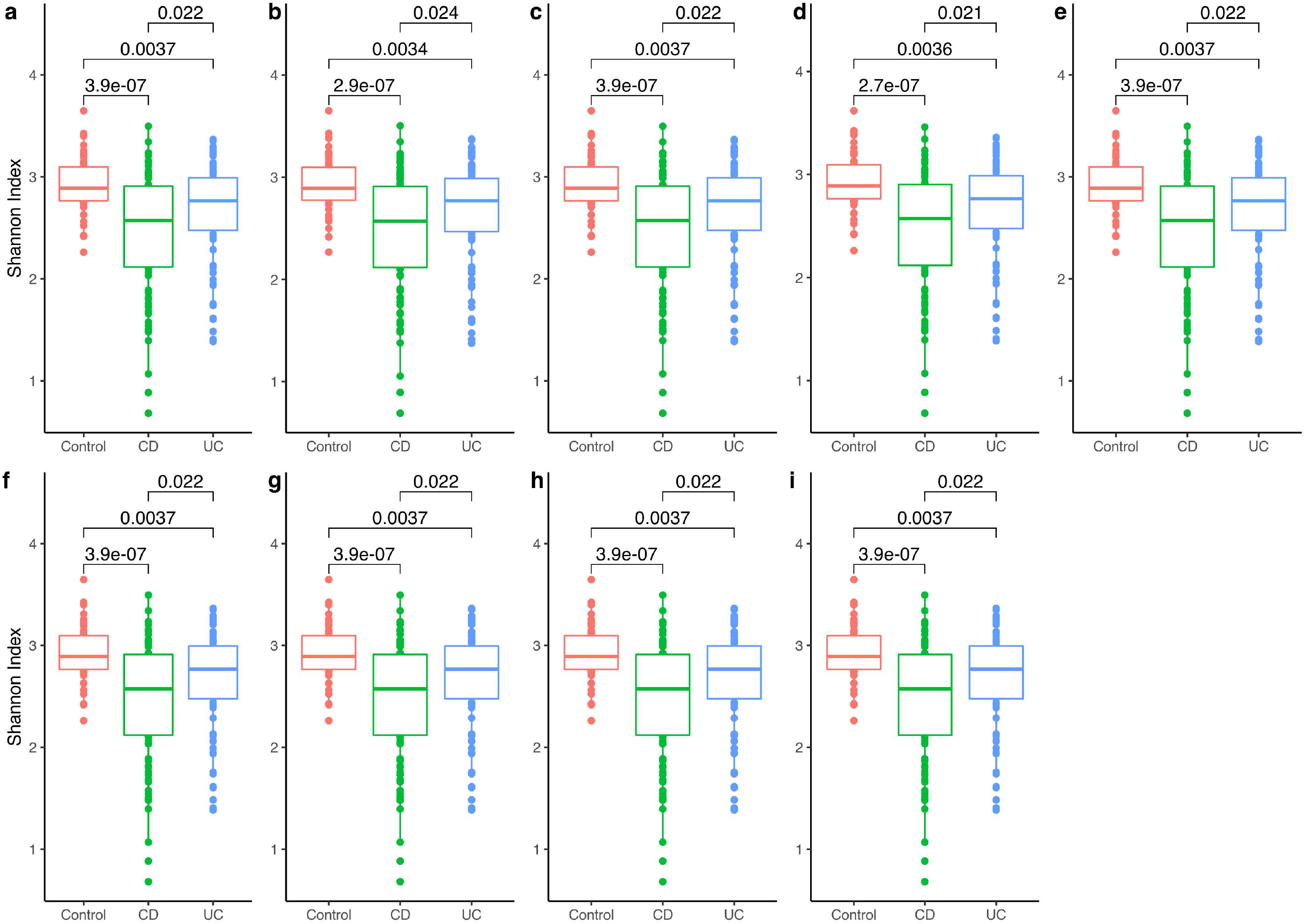
Diversity ***α*** measured by the Shannon index for different normalization methods. The statistical differences have been made through the Wilcoxon rank sum test, a) Original data; b) Rarefaction; c) ANCOM-BC; d) CSS; e) DESeq2; f) GMPR; g) TMM; h) UQ; i) Wrench. CD: Chron’s disease; CU: ulcerative colitis.

Before analyzing the diversity *α*, for the rarefaction method, it has been chosen as the cut-off value of the library in 10000 reads from the rarefaction curve obtained with the myrlin R package. In the Supplementary Material 1, a rarefaction plot from phyloseq at 10000 reads are attached.

### β diversity

This is the main parameter on which this manuscript has been focused in order to be able to evaluate the possible differences in results obtained depending on the normalization/transformation method used. For this reason, given its length, it has been considered appropriate to present the results separately for each type of statistical analysis performed.

### Species associated with pathology groups

This subsection presents the results of the alternative analysis to SIMPER that has been carried out (multipatt() function, see Materials and Methods) to infer those species that define each of the groups of the analyzed microbiome study: control group, a group with Chron’s disease (CD) and a group with ulcerative colitis (UC). Specifically, 25 species define the control group, 20 the CD group and two the UC group. Table 4 provides a list of the species selected for each diagnostic group.

**Table 4.** Species that define each pathology group: control group, Chron’s disease group (CD) and ulcerative colitis group (UC).

| Control | CD | UC |
| --- | --- | --- |
| *Coprococcus catus* | *Veillonella parvula* | *Bifidobacterium bifidum* |
| *Coprococcus comes* | *Leuconostoc citreum* | *Bifidobacterium dentium* |
| *Eubacterium hallii* | *Clostridium clostridioforme* |  |
| *Dorea formicigenerans* | *Dialister invisus* |  |
| *Subdoligranulum unclassified* | *Lachnospiraceae bacterium 2 1 58FAA* |  |
| *Roseburia hominis* | *Lachnospiraceae bacterium 5 1 57FAA* |  |
| *Ruminococcus bromii* | *Veillonella atypica* |  |
| *Dorea longicatena* | *Clostridiales bacterium 1 7 47FAA* |  |
| *Coprococcus sp ART55 1* | *Lachnospiraceae bacterium 9 1 43BFAA* |  |
| *Alistipes shahii* | *Collinsella intestinalis* |  |
| *Eubacterium ramulus* | *Blautia hansenii* |  |
| *Ruminococcus obeum* | *Pediococcus acidilactici* |  |
| *Ruminococcus sp 5 1 39BFAA* | *Coprobacillus unclassified* |  |
| *Gordonibacter pamelaeae* | *Acidaminococcus unclassified* |  |
| *Lachnospiraceae bacterium 3 1 46FAA* | *Dorea unclassified* |  |
| *Bacteroidales bacterium ph8* | *Clostridium bolteae* |  |
| *Eubacterium rectale* | *Lachnospiraceae bacterium 4 1 37FAA* |  |
| *Akkermansia muciniphila* | *Lactobacillus salivarius* |  |
| *Barnesiella intestinihominis* | *Fusobacterium nucleatum* |  |
| *Bifidobacterium catenulatum* | *Clostridium hathewayi* |  |
| *Prevotella copri* |  |  |
| *Phascolarctobacterium succinatutens* |  |  |
| *Eubacterium ventriosum* |  |  |
| *Ruminococcus lactaris* |  |  |
| *Parabacteroides goldsteinii* |  |  |

### Statistical tests

Next, Table 5 presented the results of certain statistical analyzes that have been carried out at the level of β diversity in which the Bray-Curtis distance measure has been used for the normalization methods and the Euclidean distance for the CLR *(centered log-ratio)* transformation method.

**Table 5.**
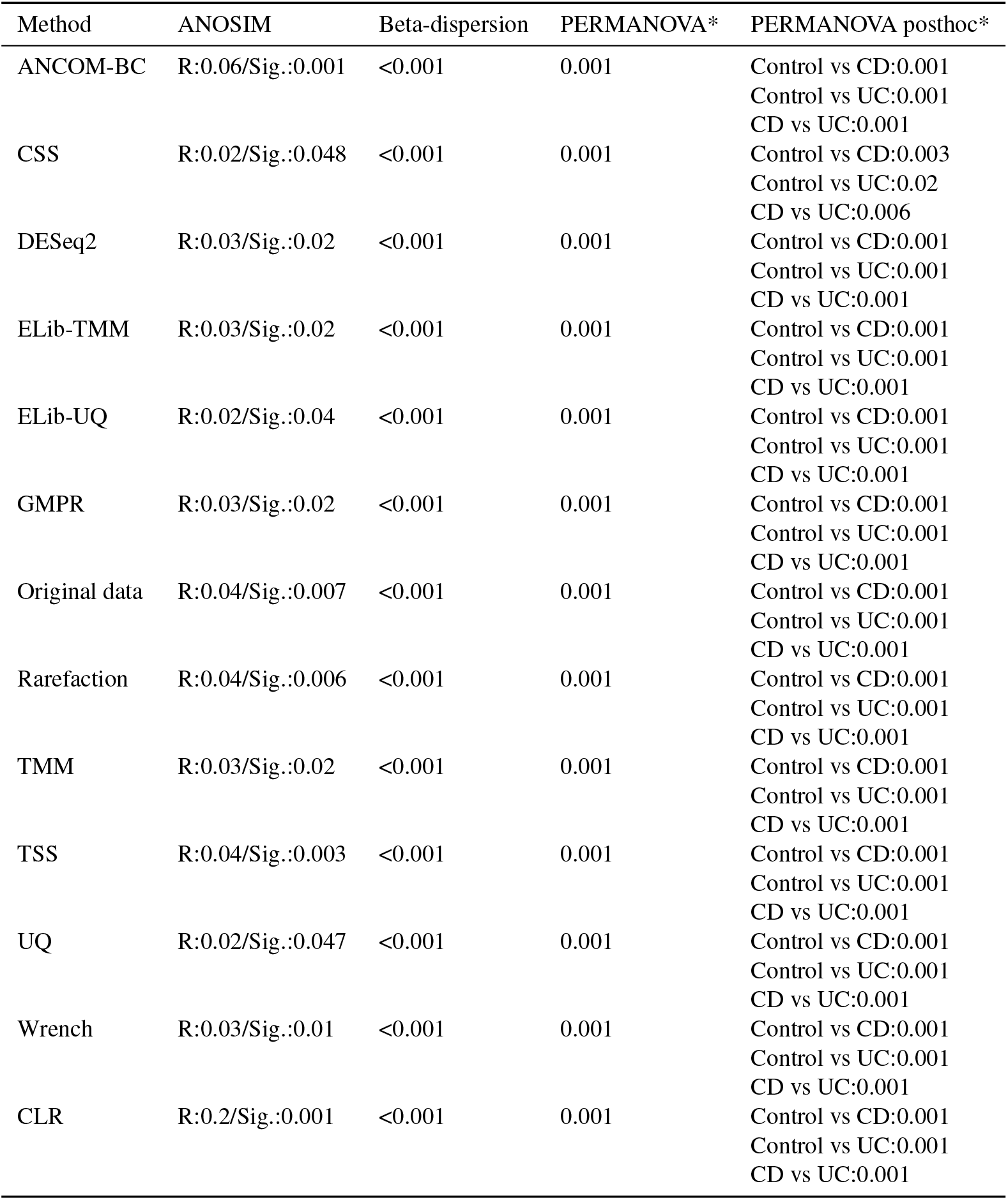
Summary of the results obtained in the statistical tests. * p-value adjusted by the false discovery rate method.

### Exploratory analysis: PCoA

The exploratory analysis of principal coordinate analysis (PCoA) has ordered the three pathology groups of the study in space in a different way depending on the normalization method studied and, in addition, the first two components explain a different percentage of variability (see Figure 5). As indicated in Materials and Methods, only the first two components are indicated. In each graph, the name of the sample that is further away from the center of the PCoA has been indicated according to the following color code: green (control group), purple (CD group), and blue (UC group).

**Figure 5.**
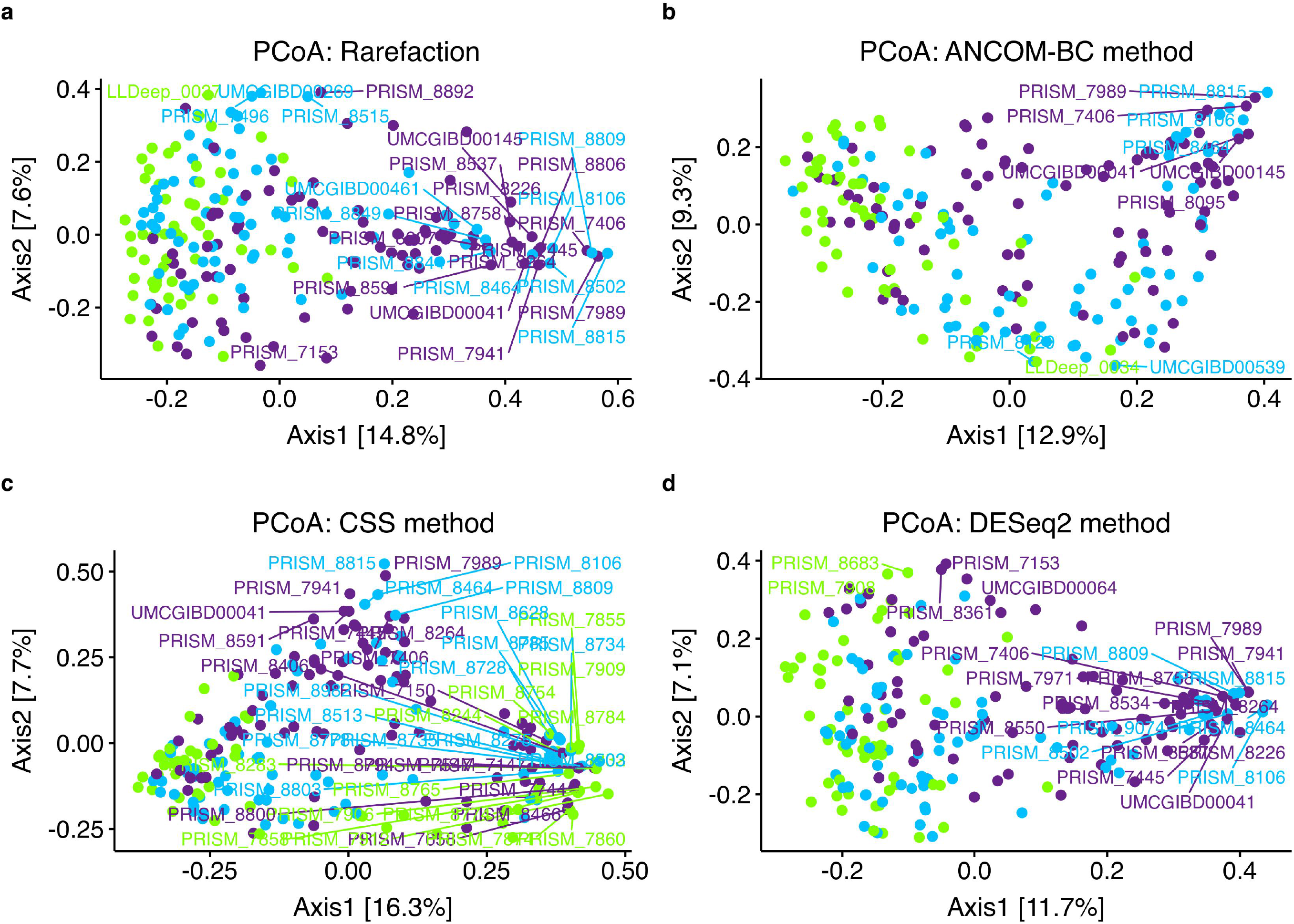
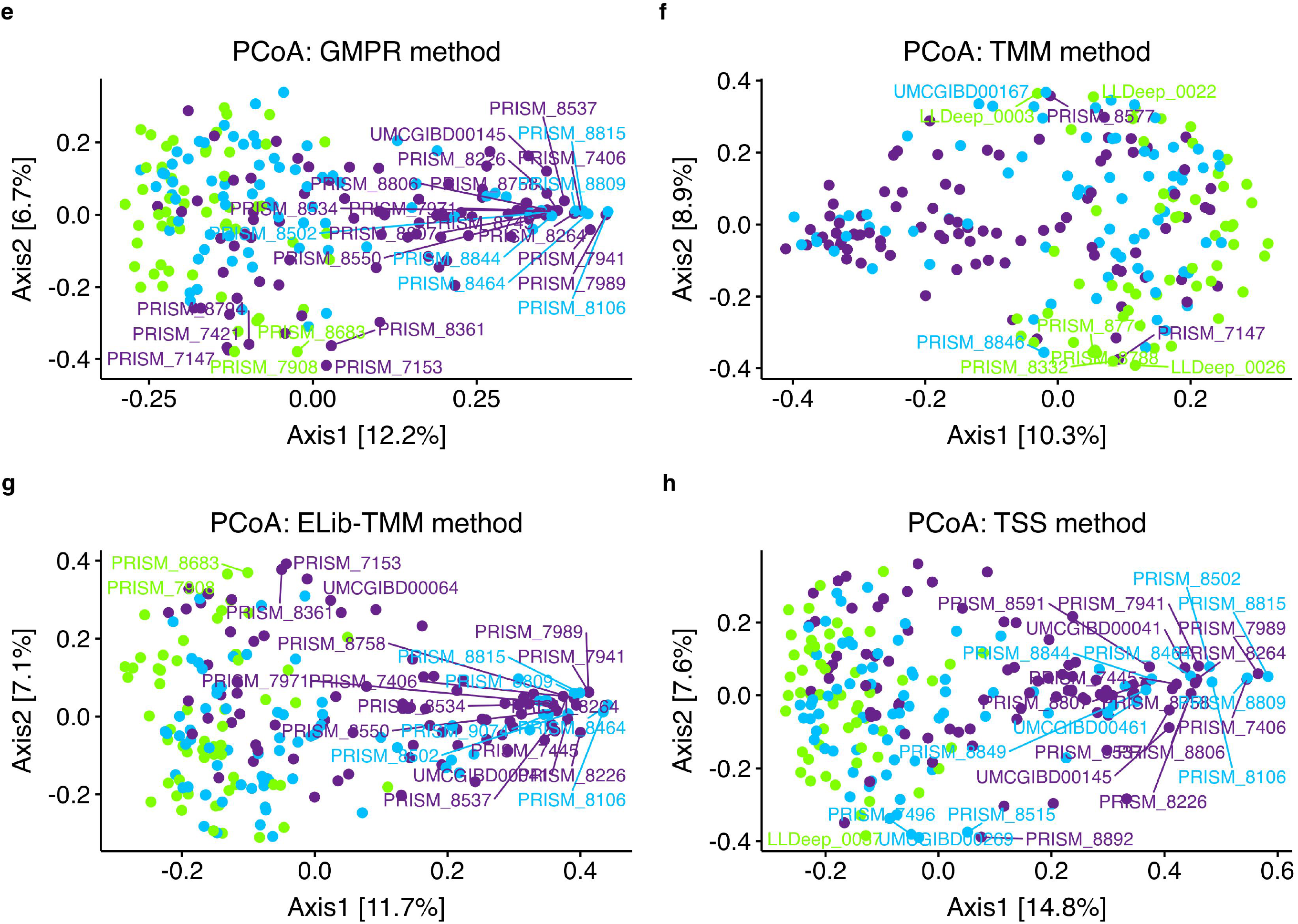
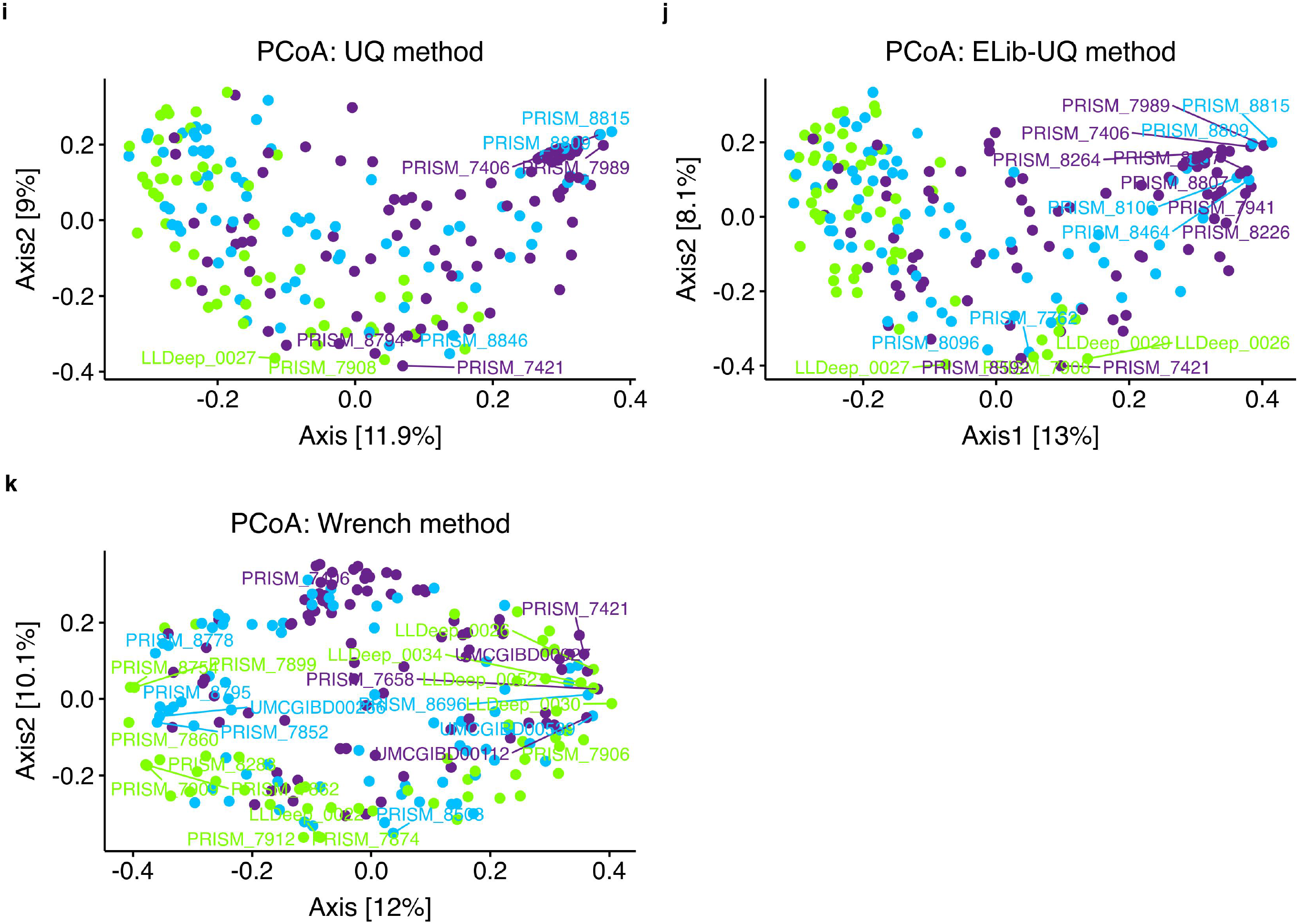
PCoA of beta diversity using the Bray-Curtis distance. The green dots are the samples from the control group; purple dots, samples from the Chron’s disease (CD) group; and, the blue dots those of the ulcerative colitis (UC) group.

The rarefaction normalization methods, TSS, DESeq2, GMPR, and ELib-TMM have presented a very similar organization in space, with the rarefaction method and TSS explaining a higher percentage of variability (22.4%).

From the PCoA obtained, we can see that the control group (green dots) is the one with a more evident cluster (the differences in species between the control samples are lower than that of the other groups). The blue color group (UC) is also quite close together but is already more dispersed than the control group. The CD group (purple color) is the most dispersed of the 3, especially along the first component (PCoA1, axis 1). In addition, it is important to point out that the two methods that explain the most variability (rarefaction and TSS) also coincide with the samples of the group UC (purple) and CD (blue) that present the composition of species further away from the rest of the samples corresponding to their group. Specifically, they are samples PRISM_8815 for the CD group and PRISM_7989 for the UC group. Finally, in the case of the CLR transformation, and as indicated in the Materials and Methods section, it was not possible to perform a PCoA but rather a PCA with the Aitchison distance (equivalent to the Euclidean distance). The PCA obtained (Figure 6) is similar to the two normalization methods that explain more variability, although the control group (green) does not appear as a group, as we can see in the TSS and rarefaction method.

**Figure 6.**
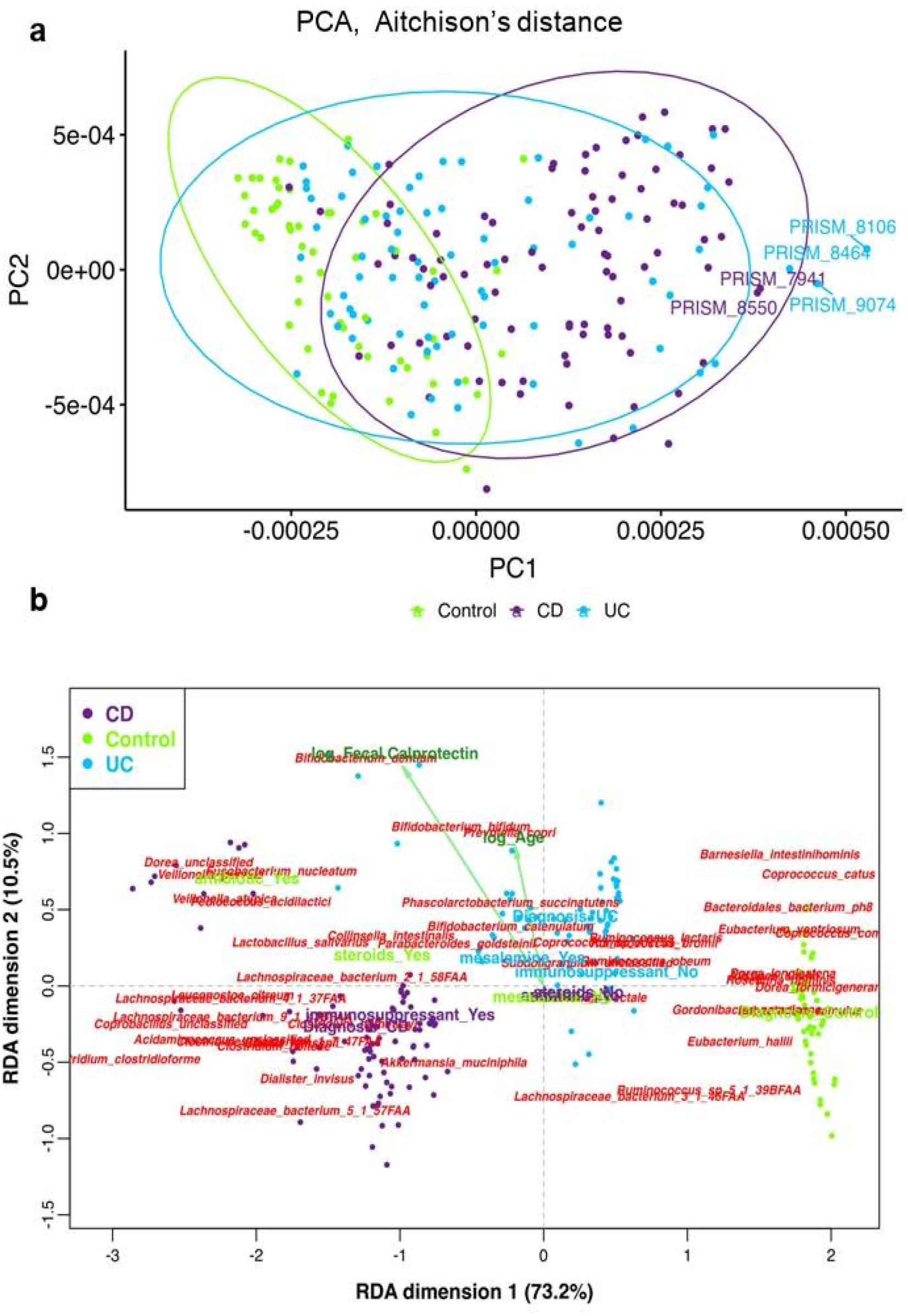
a) *β* diversity of the CLR *(centered log-ratio)* transformation method through a PCA with Aitchison distance.b) Redundant analysis with the CLR transformation method on the subset of 47 species to facilitate interpretation of the graph.

### Interpretative analysis: db-RDA

Figure 7 shows the graphs obtained from the redundancy analysis based on the Bray-Curtis distance for all the normalization methods. For the case of the “centered log-ratio” transformation, see Figure 6 since we could not use the Bray-Curtis distance, and we used another R package (easyCODA) as mentioned in Material and Methods. For all the representations, we can see that the arrangement of the species in the first two dimensions is the same among all the normalization methods, except for the TMM method (Figure 7h), which is inverted. As can be seen, what changes are the explanatory variables that the db-RDA model has automatically selected “forward” (see R script for details) and the direction of these variables in the space. From all the graphs, for example, we can conclude that the species *Ruminococcus gnavus* is related to the diagnosis of CD (Chron’s disease). However, also, according to another normalization method, it is related to taking immunosuppressants.

**Figure 7.**
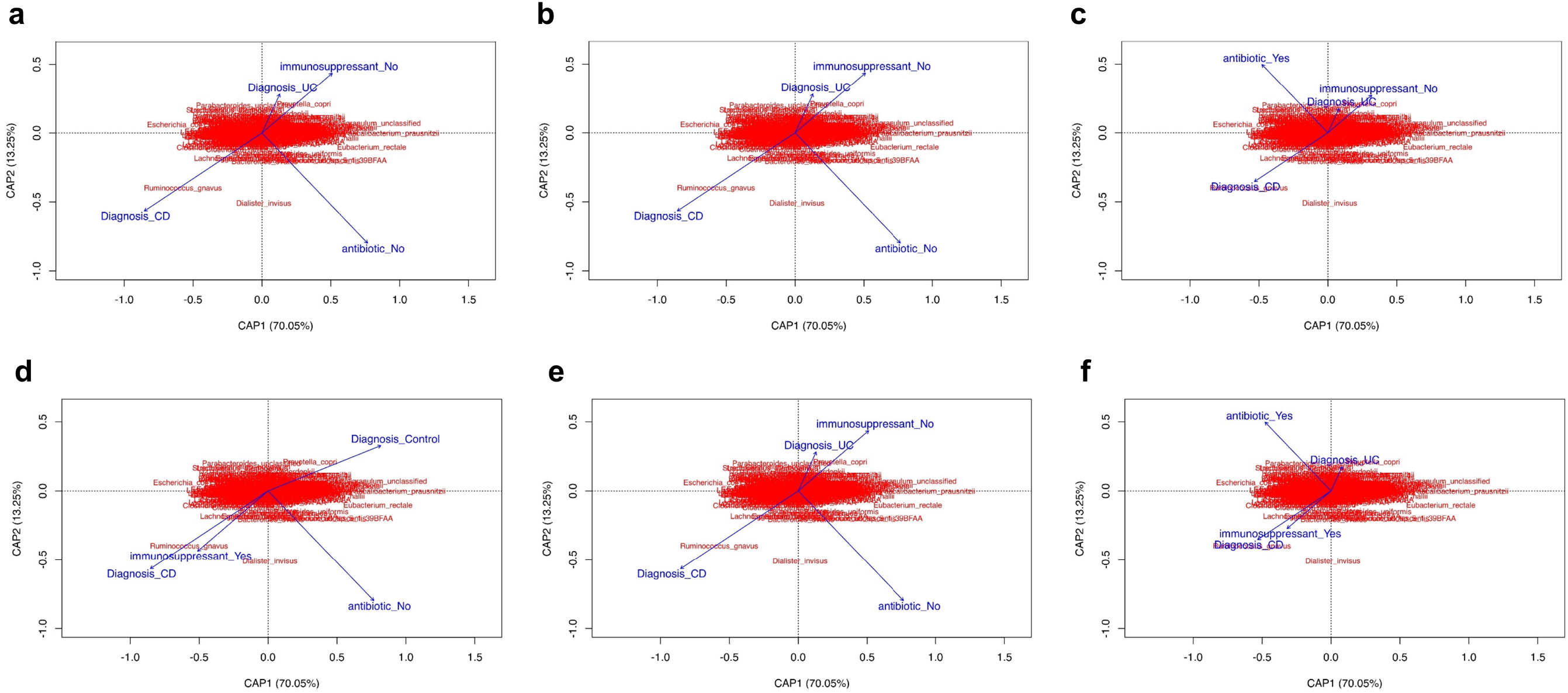
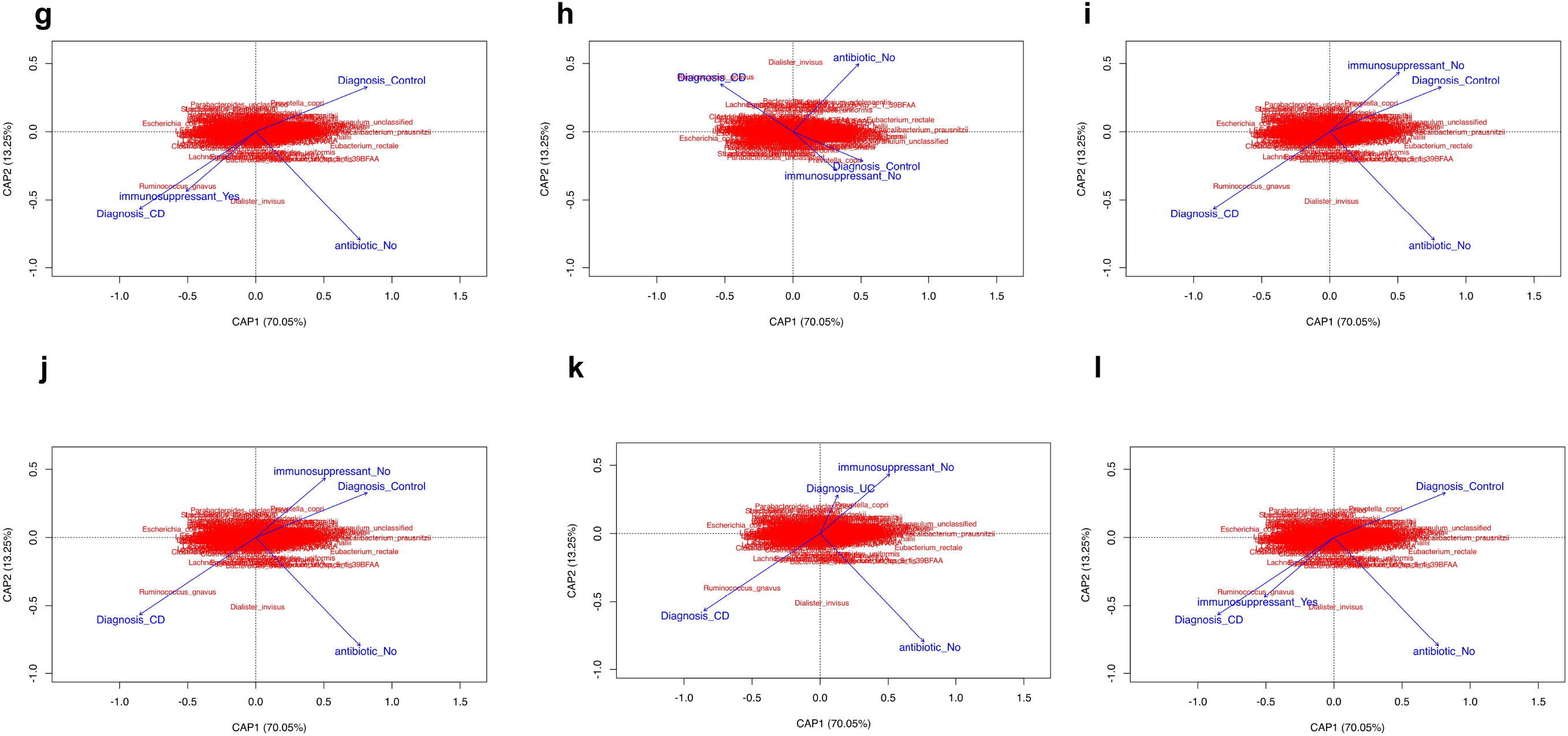
db-RDA graphs for the different normalization methods, a) ANCOM-BC; b) CSS; c) DESeq2; d) ELib-TMM; e) ELib-UQ; f) GMPR; g) original data; h) Rarefaction; i) MMT; j) TSS; k) UQ; and 1) Wrench.

### Discriminant analysis: sPLS-DA

Figure 8 shows the two main graphs obtained by each type of normalization and transformation method analyzed: the graph of the individuals and the corresponding circular correlation graph that both graphs complement each other to be able to draw conclusions. For all the methods compared, the circular correlation graph does not change, and only the groupings in the graphs of the individuals change depending on the method used. Of all the methods, the method that has explained the most variability in the first two components has been the CLR method. However, the results of the circular correlation graphs when all the species have been considered (see Supplementary Material 3) showed different species spatial organization depending on the normalization method.

**Figure 8.**
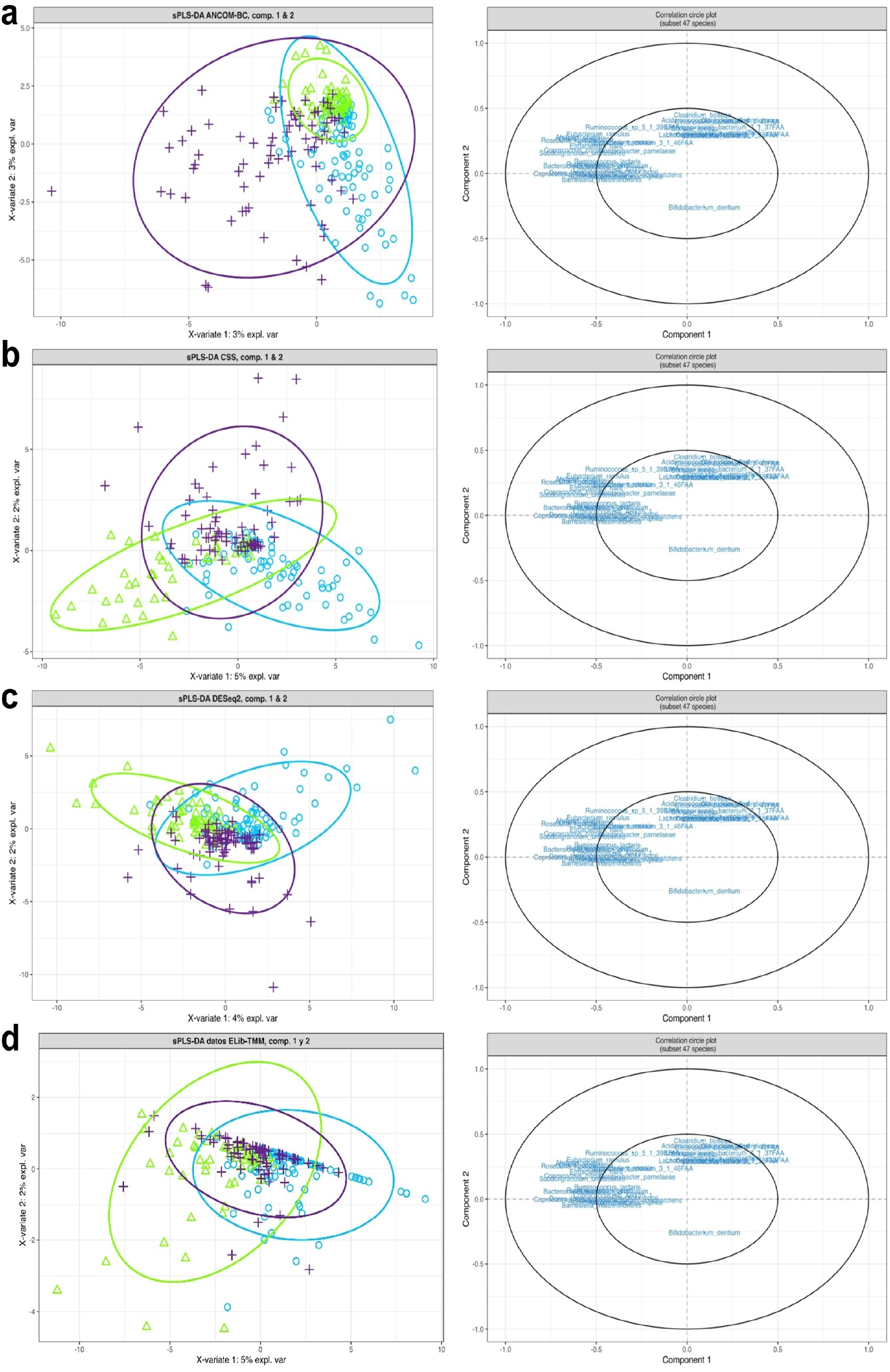
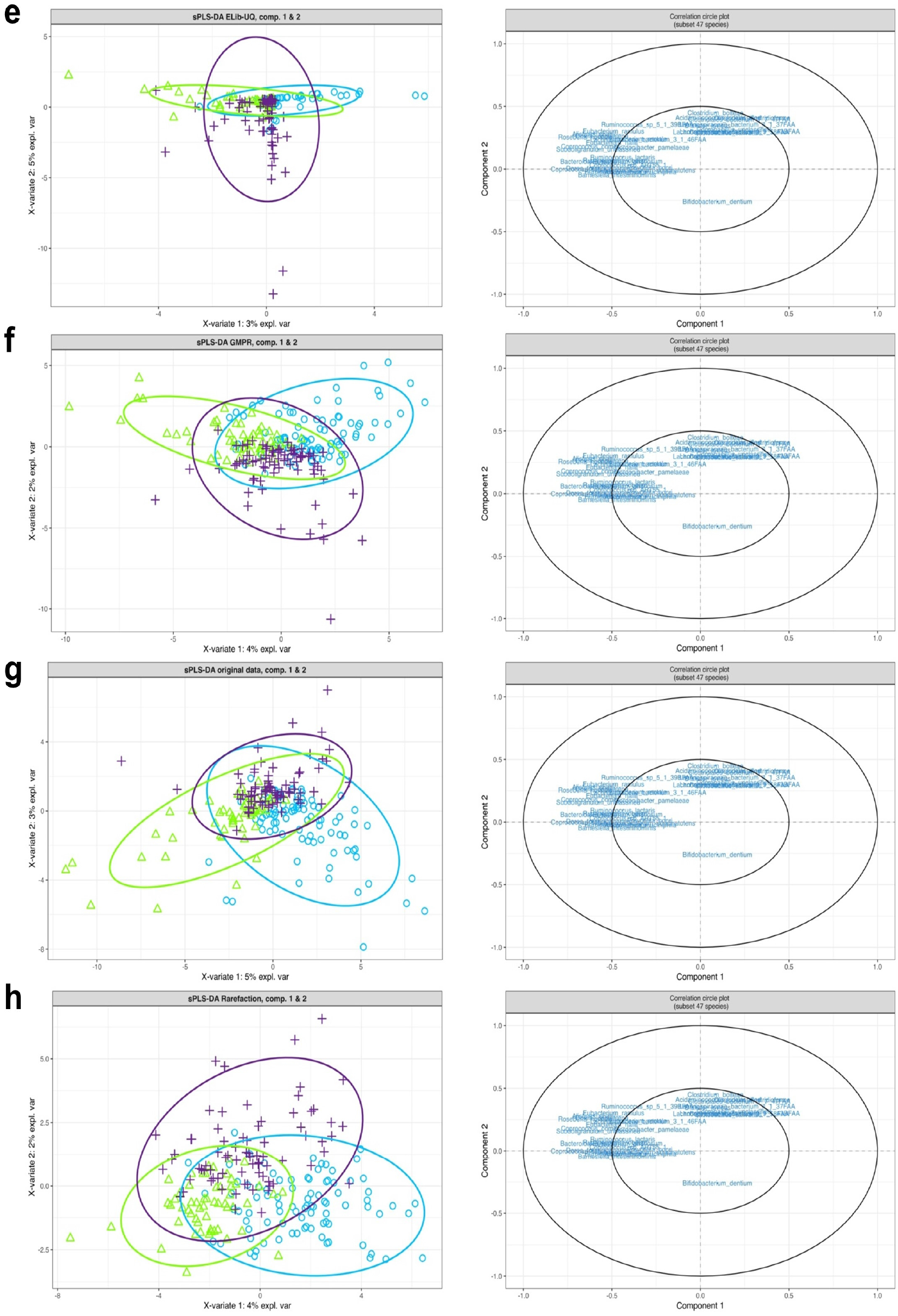
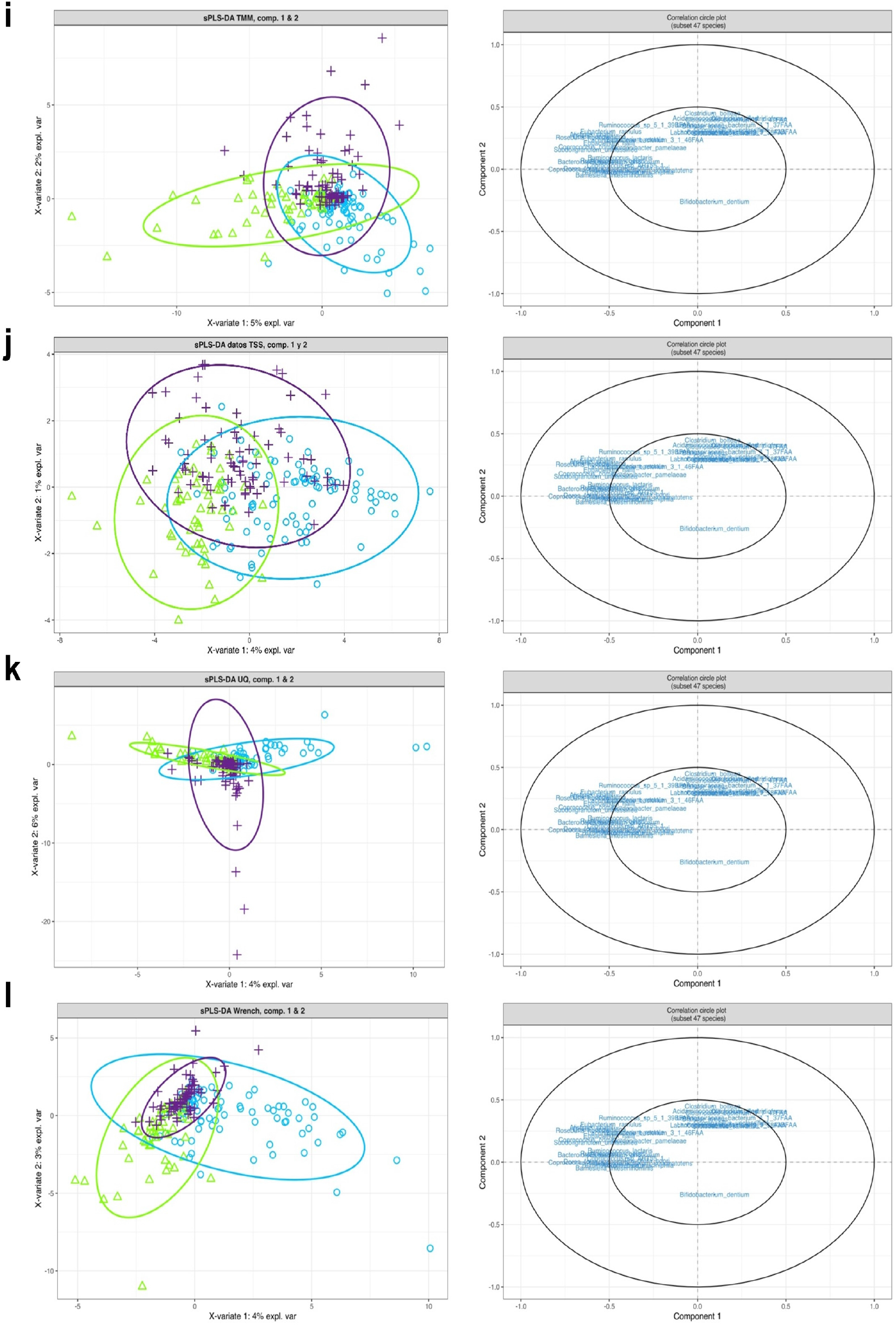
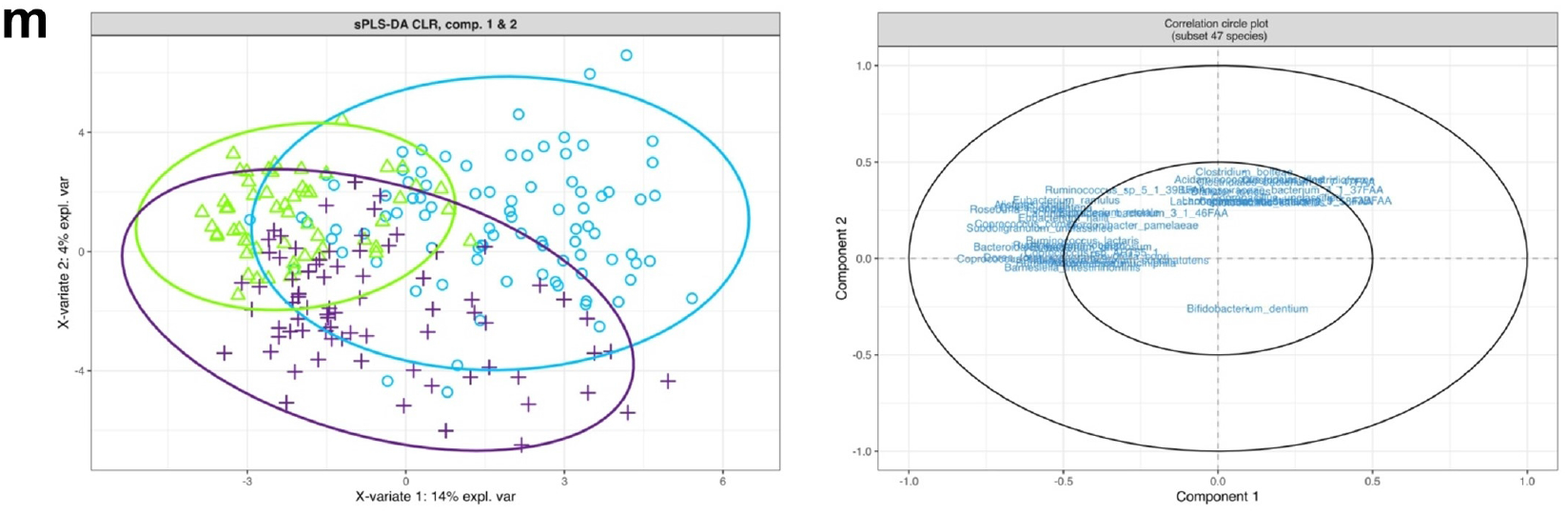
sPLS-DA plots for the 47 species subset. For all the species, see Supplementary Material 3.

In addition, as supplementary material (Supplementary Material 4), it is possible to visualize which variables have contributed the most to the first component through the plotLoadings() function of the mixOmics R package. All methods have presented the same species for each type of group. Finally, also in the Supplementary material 3, the background prediction graphs are attached, which offer an idea of the prediction for new samples in which diagnostic group they would be included.

## DISCUSSION

### Comparison between normalization methods

An adequate normalization method will eliminate (reduce as much as possible) the biases and variations introduced during the sampling and sequencing process; therefore, the normalized data will reflect biological differences. Due to its great importance, it is highly recommended to validate all possible post-normalization analyses [64].

Figures 2 and 3 have evaluated the scaling factor (and random sub-sampling for the rarefaction case) in creating a new species matrix (species rows, sample columns) between the different normalization methods evaluated in this work. Therefore, in this first analysis, a subsequent analysis of the normalized data has not been addressed, but an attempt has been made to evaluate the normalization itself.

In Figure 2, centered residuals have been evaluated on real microbiome data set following the strategy used by [33, 38] who used it to evaluate their ANCOM-BC normalization method on simulated data. From the results obtained in the real dataset, it is a bit suspicious the spectacular result obtained by the ANCOM-BC method [33, 38]. To calculate the true sample fraction (needed in ANCOM-BC method), it has been inferred through the same ANCOM-BC method (see script of R in Zenodo repository and the original script of the authors that show the functions that it is has been used to calculate it: https://github.com/FrederickHuangLin/Microbiome-Review-Code-Archive/blob/master/scripts/data_generation.R#L142. However, it is very interesting to mention that in the public dataset used in the present study, it has not found any grouping by diagnostic groups for any type of method, an undesirable aspect, and that they did detect Huang Lin’s studies for their simulated data [33, 38].

The initial idea was to reproduce figure 6 of the article by Li Chen et al. 2018 [42], but it was not possible to receive some hints from their corresponding author. However, an alternative strategy has also offered us valuable information through the coefficient of variations. In Figure 3, a box plot has been presented, taking into account the difference in the coefficient of variation in percentage, which is a standardized measure of the dispersion of the data. The GMPR method for the public dataset has presented a good aptitude in terms of data dispersion (assessed through the coefficient of variation) as occurred in [42] but the good results for the TMM and Wrench methods should also be highlighted, the latter not evaluated in [42].

### *α* diversity

The Shannon index has been used to compare the *α* diversity between the different normalization methods (Figure 4). As indicated in the Materials and Methods section, for its calculation, it is necessary (as is logical) that the counts of the species be 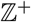. Consequently, prior to calculating the diversity index, an upward rounding was performed once the scaling factor was applied because we had decimal values. In addition, for those methods that normalized the matrix of species in relative abundances between 0 and 1 (TSS, ELib-TMM and ELib-UQ), and given that for each species of the 201 in the dataset they had a relative abundance <0.5, it was rounded to 0. Therefore, its Shannon index could not be calculated, so it does not appear in Figure 4.

No differences were observed in determining the Shannon diversity index between the different normalization methods considered. To the best of my knowledge, it has not found any literature that has evaluated normalization methods against *α* diversity. Only, a comment from the developer of the GMPR method on GitHub said that it would not affect diversity *α* https://github.com/jchen1981/GMPR/issues/2. Finally, mention that the diversity *α* for the CLR data has not been studied since it is an analysis that has not been considered [11, 59] until very recently with the recent appearance of the R coda4microbiome [65], which allows the calculation of the Shannon index but not other biodiversity indices, at least at present.

### Statistical tests related to *β* diversity

Table 5 shows the p-values for the similarity analysis tests (ANOSIM), beta-dispersion (equivalent to Levene’s test but for multivariate analysis), and PERMANOVA. The beta dispersion and PERMANOVA have been practically identical for all the normalization and transformation methods, demonstrating that the normalization/transformation method does not affect this type of analysis. However, for ANOSIM, some nuances have been obtained that it is considered appropriate to highlight briefly. ANOSIM is a non-parametric method that tests the hypothesis that there are no differences between two or more groups of samples based on the permutation test of similarities between and within groups [66]. That is, it compares the variation in the abundance and composition of species between samples taking into account a grouping factor (in our case, the patient’s Diagnosis). The null hypothesis is that there are no differences between members of the treatment groups (patient Diagnosis). In addition, to correctly interpret its results, it must consider the two values it gives us: R and significance. First, it is necessary to check that the significance value is less than 0.05. Once checked, the value of R is checked. If it is less than 0.2, it means that the chosen grouping factor (Diagnosis) has a small effect in explaining the difference between the species: http://www.researchgate.net/post/Can_anyone_help_me_in_understanding_and_clearly_interpreting_ANOSIM_Analysis_of_Similarityand_SIMPER_Similarity_percentage_analysisresults.

Going back to Table 5, we can see that all the methods turned out to be significant except almost for the CSS and UQ method. Regarding the value of R, all have presented a very small value of R, except for the CLR transformation method on the border (0.2). In summary, except for the CLR method, we can conclude that Diagnosis is not a factor variable essential to explain the difference in species presented by the three diagnostic groups considered in the analyzed study. To the best of our knowledge, it has not found any bibliographic reference that has compared ANOSIM between different normalization methods. In relation to PERMANOVA, it is noteworthy to mention that Weiss et al. 2017 [64] detected differences depending on the normalization method used and using several public datasets.

### Multivariate analysis of *β* diversity

As indicated in the Materials and Methods section, one method was performed for each large group of multivariate analyses: PCoA (exploratory analysis), db-RDA (interpretative analysis) and sPLS-DA (discriminative analysis). In all of them, the Bray-Curtis distance has been used (except for the Euclidean for the CLR method) since it is the distance par excellence in disciplines such as Ecology and the analysis of the microbiota, since, for example, Bray-Curtis gives us a better idea of the dissimilarity of the species between samples compared to the Euclidean distance, since with Bray-Curtis the maximum distance is obtained when the samples that are compared do not have species in common, among many other aspects commented on in several articles of Carlo Ricotta [67, 68].

In addition, it is very important to highlight that both the *α* and *β* diversity analyzes have traditionally been calculated (at least in Ecology and Microbiology) based on relative abundances (TSS method), but also by the rarefaction method [28, 64]. From an Ecological (and Microbiological) point of view, the main reason for using relative abundances of species rather than absolute abundances for the calculation of functional dissimilarity is that ecologists (microbiologists) are often interested in exploring how species changes the ecological strategies or evolutionary pathways of species among samples of each diagnostic type (i.e., how functional traits change and phylogenetic characteristics are proportionally distributed among species), regardless of the absolute abundances of species in each parcel (sampling unit) [68]. However, the disciples of the CODA school created by the statistician Aitchison defend the non-use of the Bray-Curtis metrics distance and also that the Bray-Curtis distance can be used for the original counts (not only for relative counts) http://www.econ.upf.edu/~michael/stanford/; https://www.youtube.com/watch?v=c7VUrViGmQU.

The first multivariate analysis that was carried out was the exploratory analysis using principal coordinate analysis (PCoA). Principal component analysis (PCA) establishes the conserved distance between two objects: the Euclidean distance. If it is desired to order the objects based on another distance measure (for example, the Bray-Curtis distance) then PCoA is the method of choice. PCoA provides a Euclidean representation of a set of objects whose relationships are measured by any user-chosen measure of similarity or distance. As in the case of PCA, PCoA produces a set of orthogonal axes whose importance is measured by the eigenvalues (*eigen values*) [53]. As expected from the discussion above, those methods based on rarefaction and relative abundances (TSS in particular) have presented an identical spatial arrangement with a higher % of variability explained by the first two components. Furthermore, normalization methods have been shown to modify beta-diversity in PCoA representation, at least in our analyzed real data, which contradicts the comment by the developer of the GMPR method in a GitHub thread https://github.com/jchen1981/GMPR/issues/2

The next multivariate analysis performed on the public data analyzed was the Bray-Curtis distance-based redundancy analysis (db-RDA). The db-RDA is an ordering method similar to redundancy analysis (RDA), but that allows the use of non-Euclidean dissimilarity indices (Bray-Curtis, for example). Despite this non-Euclidean feature, the db-RDA analysis is strictly linear, and metric [53]. The db-RDA (and the RDA as well) is an extension of multiple regression, which models the effect of an explanatory matrix ***X*** (*nxp*) on a response matrix ***Y*** (*nxm*). The difference here is that we can model the effect of an explanatory matrix on a response matrix rather than a single response variable. Therefore, the db-RDA (RDA) allows us to model the effect of medical variables (consumption of antibiotics, immunosuppressants, mesalamine…, presence of occult blood in faeces) in the entire population studied, not just in a single sample. This is achieved by sorting ***Y*** to obtain sort axes that are linear combinations of the variables in ***X***, which ***X*** is the species matrix [53], http://r.qcbs.ca/workshop10/book-en/redundancy-analysis.html. It will not expand further on this method, but we will add the concept of constrained *(constrained)* and unconstrained proportions that appear during the db-RDA parsing. The constrained proportion refers to the variance of ***Y*** explained by ***X***, while the unrestricted proportion refers to the unexplained variance in ***Y***. All the values obtained by each method can be consulted in detail in the R scripts.

As it has been commented in the Results section for Figure 7, no differences have been observed in the arrangement of the species, but yes in the direction of the explanatory variables. However, practically all the species appear superimposed on the axis (0,0) of the graph, which does not allow any interpretation beyond separated species. The most extreme case has been for the species *Ruminococcus gnavus* related to Diagnosis CD that was not identified by the alternative SIMPER strategy followed (see Table 4). Apart from the biplot presented in Figure 7, a triplot has also been made (contains sample information) with the use of the ggord() function, and its graphs have been included as Supplementary Material 2.

For the case of the CLR transformation method, an RDA has been performed since the Euclidean distance has been used (see Figure 6b). As can be seen, the RDA triplot of the CLR method allowed us to make many more groupings of species per diagnostic group and also relate them to more explanatory variables. Therefore, in summary, the CLR method has turned out to be more informative than any other method in the interpretive analysis through redundancy analysis, and it could be more advisable to follow an approximation of the microbiome data as compositional data as sustented by several authors [11, 29].

The third and last multivariate analysis was the sPLS-DA as a discriminative analysis technique [53]. sPLS-DA is an extension of the sPLS method, a regression technique initially applied to chemometrics but was found to be useful on omics data. The sPLS adds *sparsity* into the PLS with a Lasso penalty combined with an SVD computation. For complete detail, see [69]. Although the PLS method was primarily designed for regression problems, it works well on classification problems. SPLS-DA performs the selection of variables and classification in a single step and is a machine learning technique because it will allow us to make predictions and find a microbiological species signature for each diagnostic group [70, 71].

As commented in the results section for Figure 8, no differences were observed between the normalization methods and the circular graphic’s CLR transformation method. However, a great difference has been detected when we consider the graph of the individuals. For example, we have the species *Bifidobacterium dentium* that in Table 4 we found to be related to the ulcerative colitis (UC) group, and if we look again at Figure 8, we can infer that for the UQ and CLR methods they would be inside the UC ellipse (fucsia color). However, when we consider the background graphs (attached as Supplementary Material 3), we see that apart from CLR and UQ that would present the same prediction commented above for *Bifidobacterium dentium*, the DESeq2 method, ELib-UQ and GMPR would also match. Finally, it is important to highlight the results of the loading plots (Supplementary Material 4 that indicate the contribution of each variable for each component). The loading graphs for the first component are attached. For all normalization methods, for the first component, indicated the same signature of species for each diagnostic group. The only study found that compared normalization methods with sPLS-DA found no difference between the two methods compared: CSS vs TSS+CLR [71]. The study [71], also like the present work, used public data.

## CONCLUSIONS

A composition describes the parts of a whole quantitatively. The compositional information it contains is considered to reside in the ratios between any of the parties considered. Microbiome data are compositional data that are also characterized, like other omics disciplines, by presenting a high percentage of zeros (to denote, for example, that a certain taxon has not been detected for a specific sample) and a high dispersion in the values of taxa counts. For this reason, since the decade of the 70s, and thanks to the work of several scientists in the discipline of Ecology and Statistics, it has allowed the appearance of several methods of normalization and transformation of taxa counts, which a posteriori, have been applied to the statistical analysis of various omics, such as the case of the microbiome.

The present work has attempted to present and compare the vast majority of available normalization methods (11 methods) for the microbiome analysis in any discipline (soil ecology, clinical medicine…) and emphasize its main alternative: centered log-ratio, CLR, from CODA school disciples.

To achieve the main objective discussed in the previous section, public results of a microbiome study conducted in the United States have been used instead of simulated data (a common strategy detected in the literature consulted). Analyzes have been carried out to compare the output obtained between the different normalization methods and how each normalization and transformation method affects the *α* and *β* diversity, which are rarely addressed in the scientific literature.

The GMPR (geometric mean of pairwise ratios) normalization method presented the best results regarding dispersion of the new matrix obtained after being scaled. For the case of *α* diversity, no differences were detected among the normalization method compared. In terms of *β* diversity, the redundancy analysis as well as the sPLS-DA analysis have allowed us to detect meaningful differences between the normalization methods, being the CLR transformation method the most informative, allowing us to make more predictions. It is important to emphasize that the CLR method and the UQ normalization method have been the only ones that have allowed us to make predictions from the sPLS-DA analysis, so their use could be recommended for other real datasets.

## ACKNOWLEDGEMENTS

DB-C would like to thank the help and critical reading of this work by the ecologist and statistician Dr. Rosana Ferrero and Prof. Juan Luis López from Máxima Formación

