## Supplementary Document 1 for "Alpha and beta-diversities performance comparison between different normalization methods and centered log-ratio transformation in a microbiome public dataset"

**Supplementary Material 1.**

Rarefaction curve with the library size of 10000 reads. A plateau phase is obainted and no sample is lost.

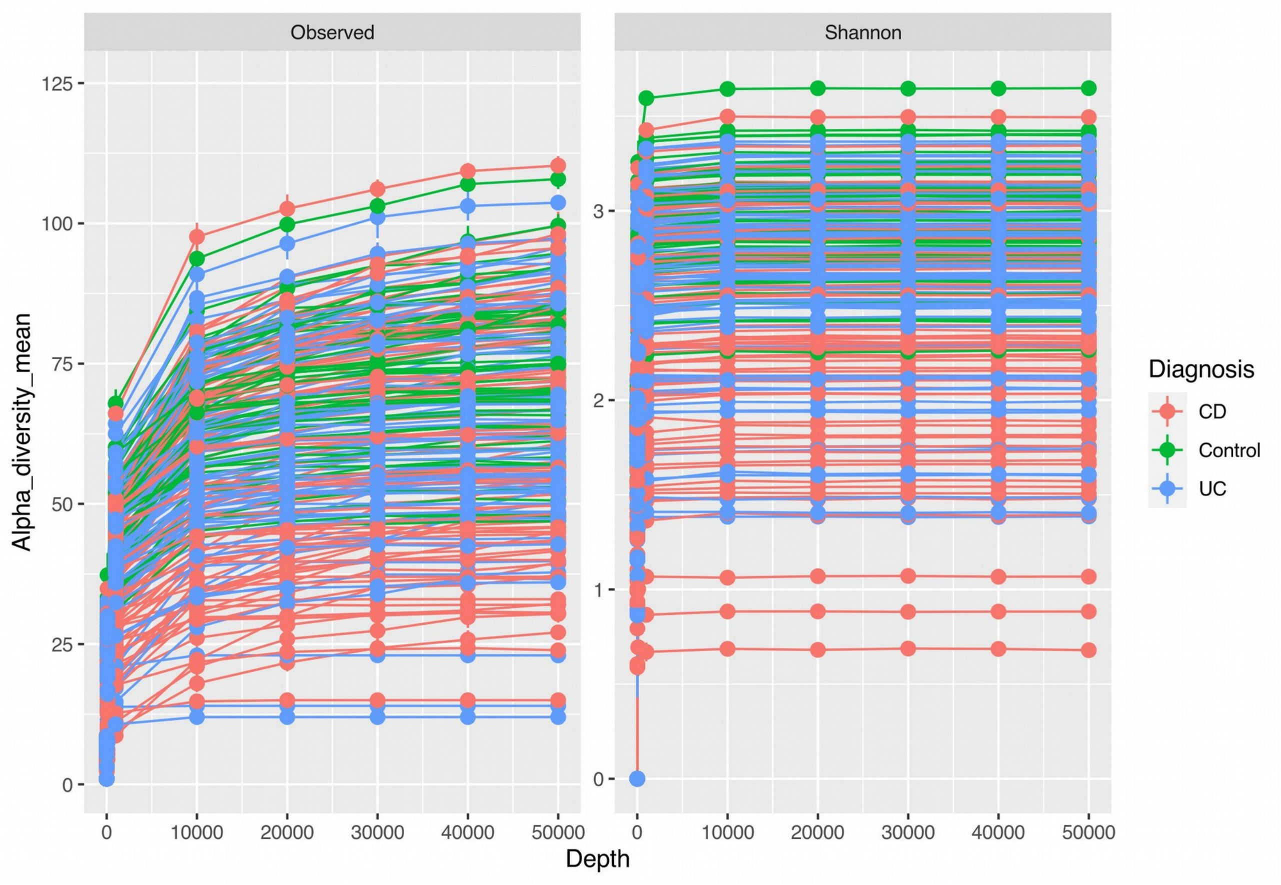
