## Supplementary figures and images for "Alpha and beta-diversities performance comparison between different normalization methods and centered log-ratio transformation in a microbiome public dataset"

### Supplementary Document 2

**Supplementary Material 2**

db-RDA plotted through ggord() R function.

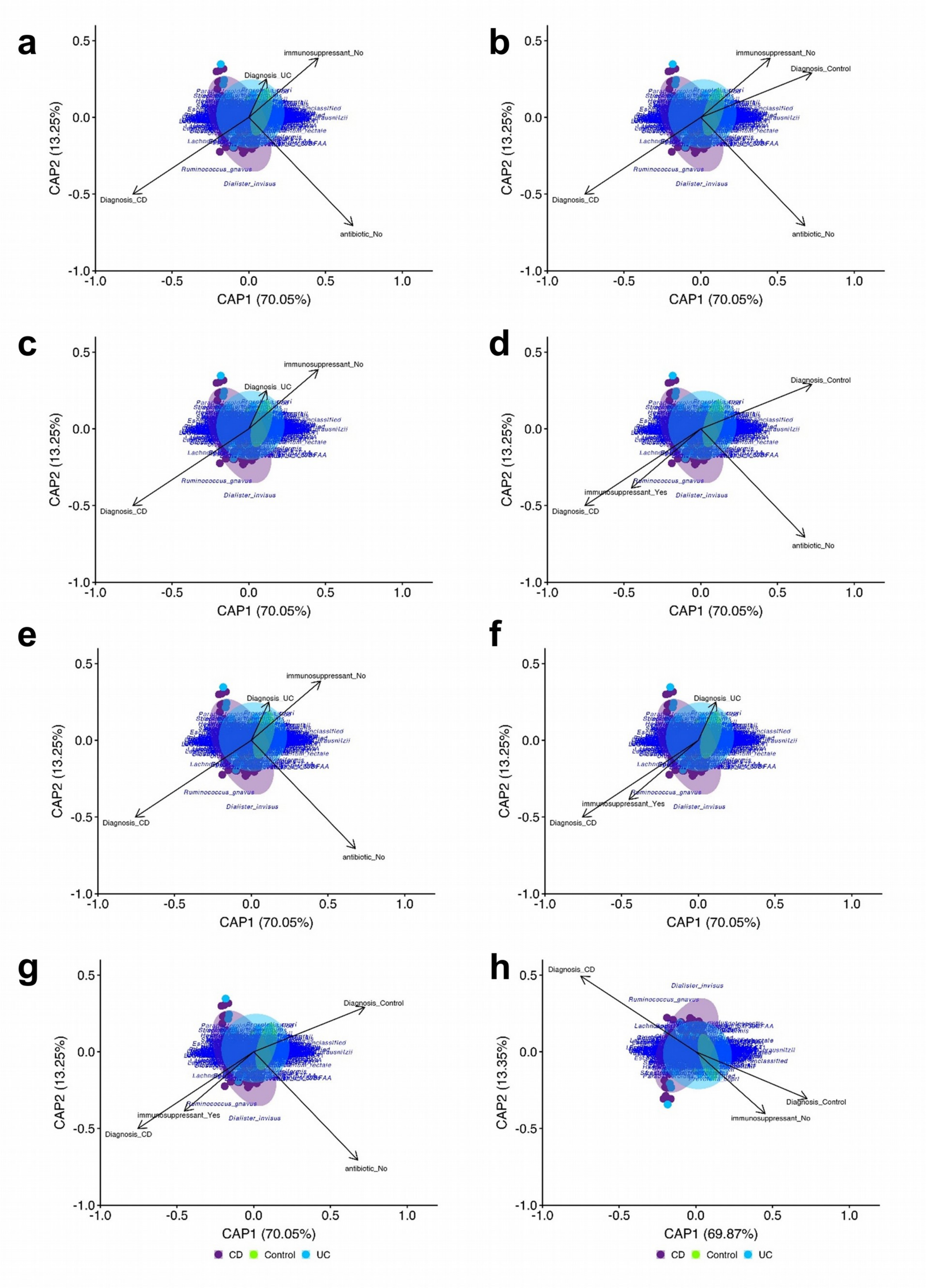

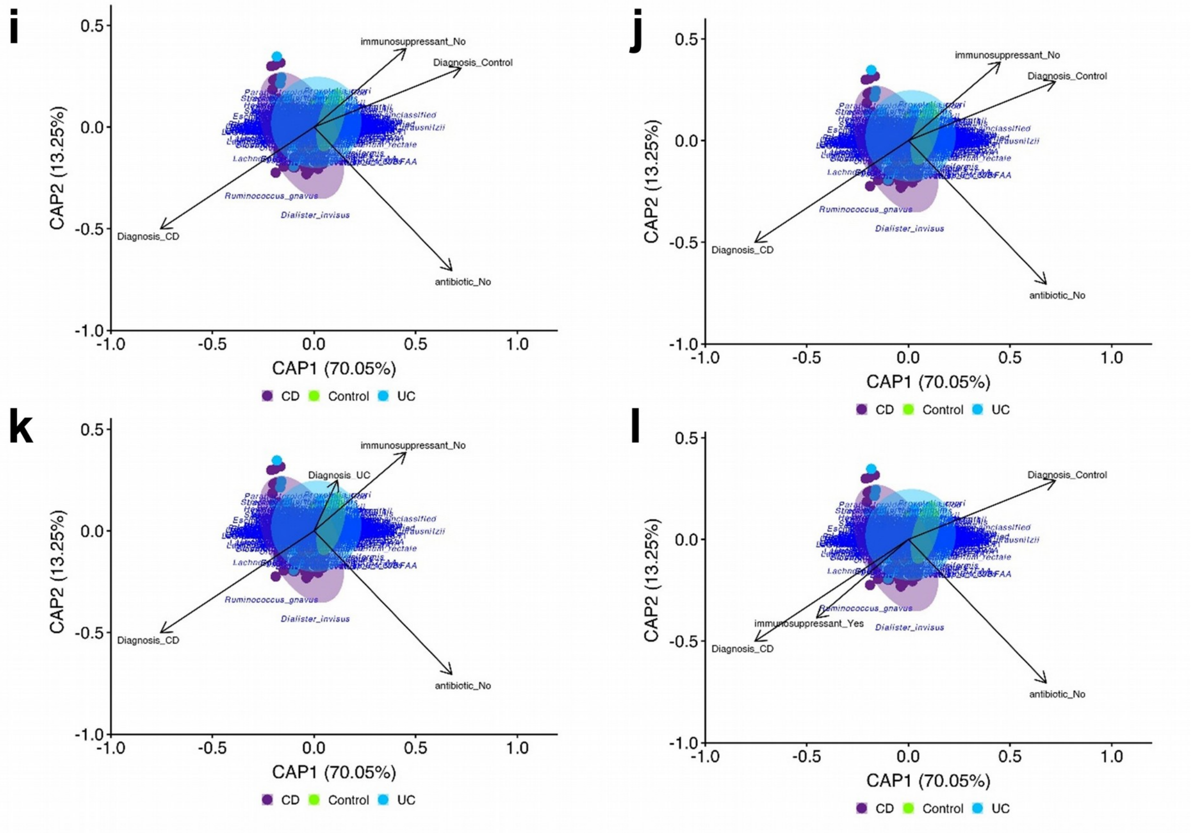
