## Supplementary Document 3 for "Alpha and beta-diversities performance comparison between different normalization methods and centered log-ratio transformation in a microbiome public dataset"

**Supplementary material 3**

sPLS-DA background prediction plots and their correlation circle plot with all species.

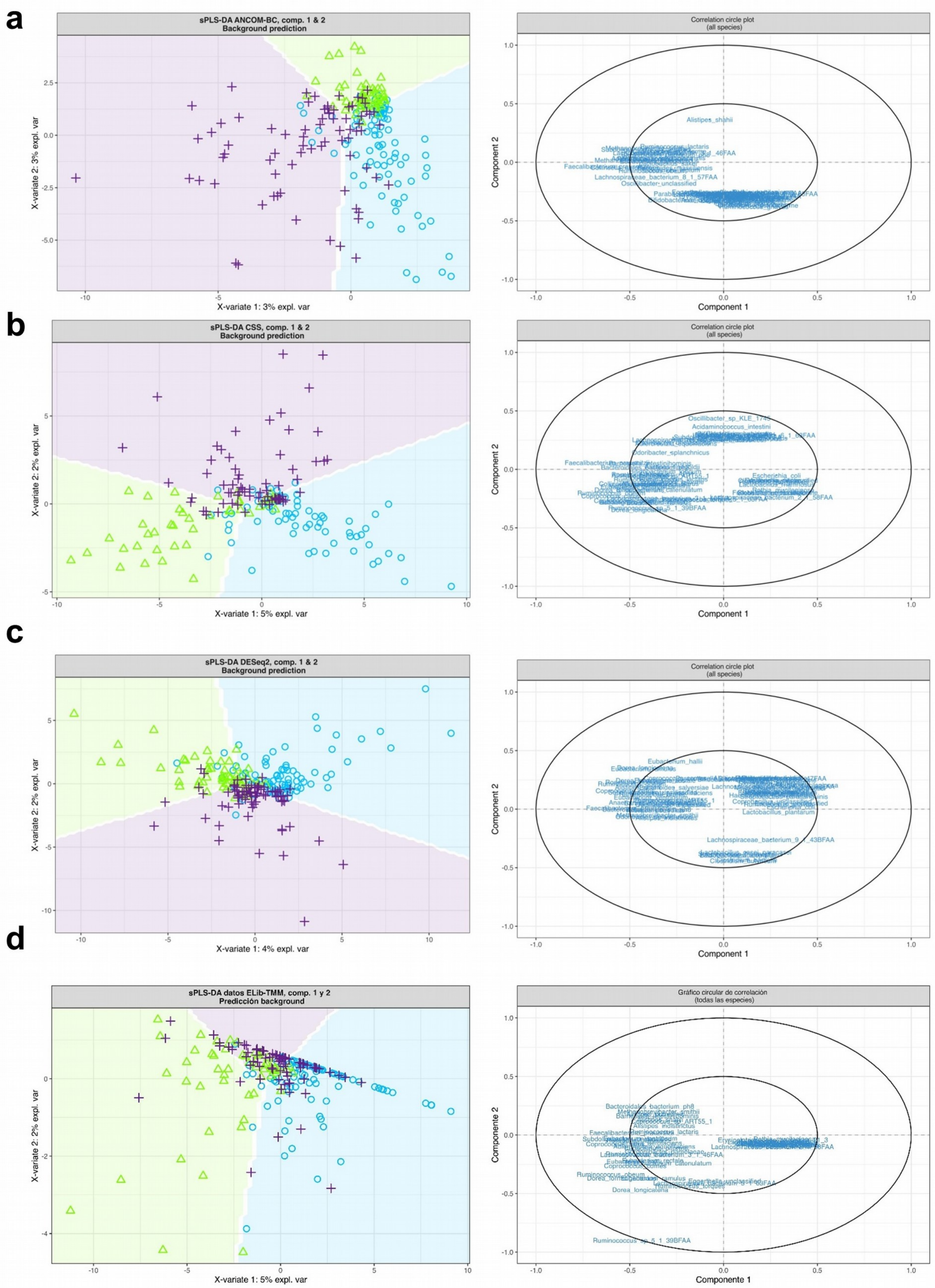
a
sPLS-DA ANCOM-BC, comp. 1 & 2
Background prediction
X-variate 1: 3% expl. var
X-variate 2: 3% expl. var
Correlation circle plot
(all species)
Component 1
Component 2
Alistipes_shahii
Lachnospiraceae_bacterium_6_1_57FAA
b
sPLS-DA CSS, comp. 1 & 2
Background prediction
X-variate 1: 5% expl. var
X-variate 2: 2% expl. var
Correlation circle plot
(all species)
Component 1
Component 2
Oscillibacter_sp_KLE_1745
Acidaminococcus_intestini
Odoribacter_splanchnicus
Escherichia_coli
c
sPLS-DA DESeq2, comp. 1 & 2
Background prediction
X-variate 1: 4% expl. var
X-variate 2: 2% expl. var
Correlation circle plot
(all species)
Component 1
Component 2
Eubacterium_hallii
Lactobacillus_plantarum
Lachnospiraceae_bacterium_9_1_43BFAA
d
sPLS-DA datos ELib-TMM, comp. 1 y 2
Predicción background
X-variate 1: 5% expl. var
X-variate 2: 2% expl. var
Gráfico circular de correlación
(todas las especies)
Componente 1
Componente 2
Bacteroidales_bacterium_ph8
Alistipes_indistinctus
Ruminococcus_obeum
Dorea_longicatena
Ruminococcus_sp_5_1_39BFAA

e

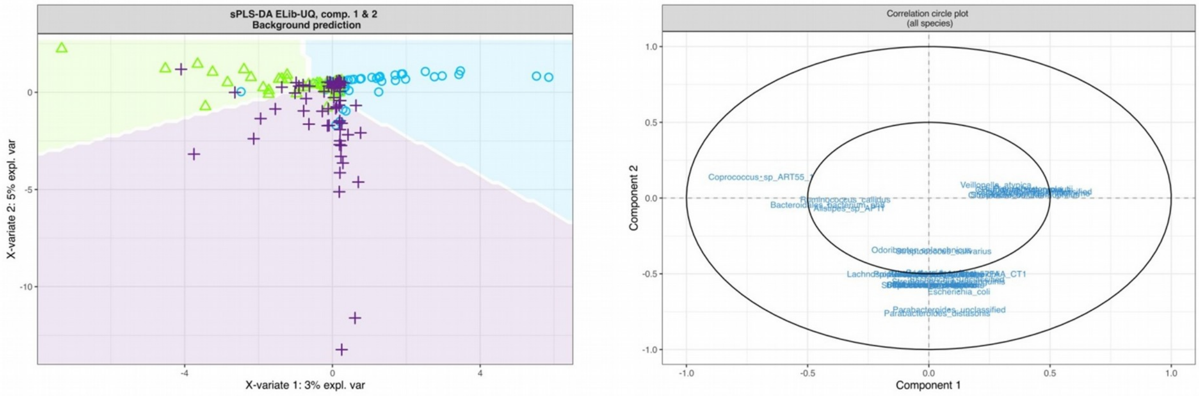
sPLS-DA ELib-UQ, comp. 1 & 2
Background prediction
X-variate 1: 3% expl. var
X-variate 2: 5% expl. var
Correlation circle plot
(all species)
Component 1
Component 2

f

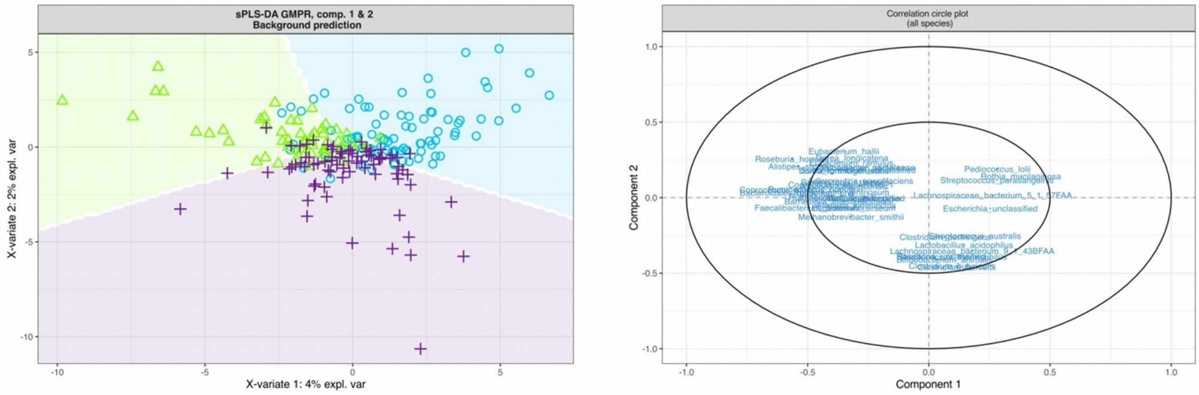
sPLS-DA GMPR, comp. 1 & 2
Background prediction
X-variate 1: 4% expl. var
X-variate 2: 2% expl. var
Correlation circle plot
(all species)
Component 1
Component 2

g

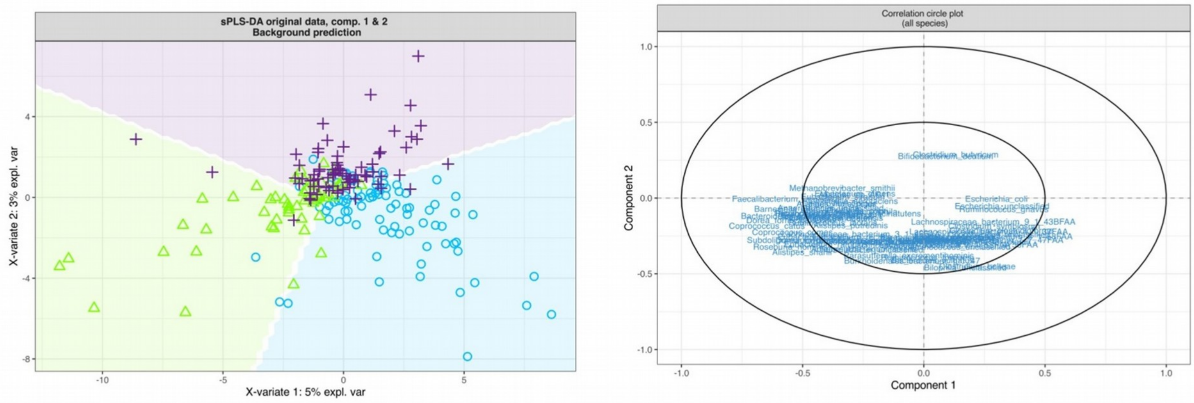
sPLS-DA original data, comp. 1 & 2
Background prediction
X-variate 1: 5% expl. var
X-variate 2: 3% expl. var
Correlation circle plot
(all species)
Component 1
Component 2

h

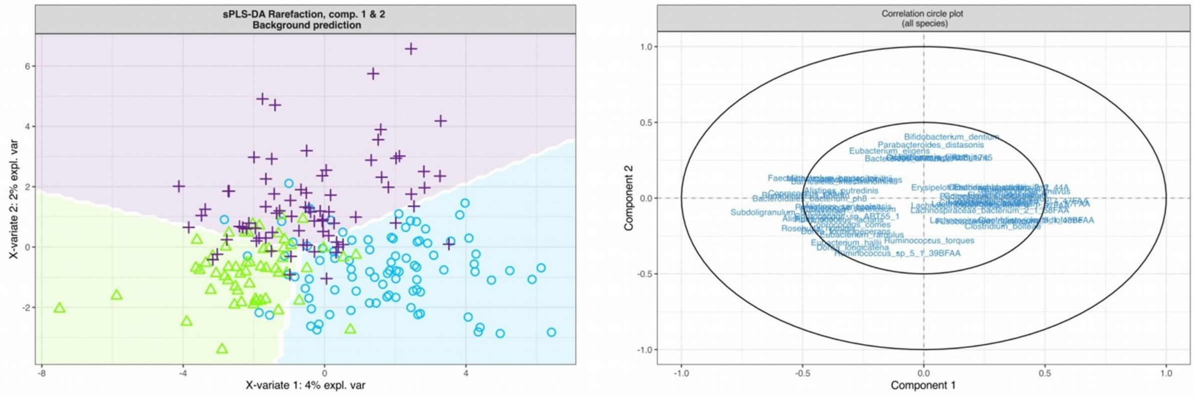
sPLS-DA Rarefaction, comp. 1 & 2
Background prediction
X-variate 1: 4% expl. var
X-variate 2: 2% expl. var
Correlation circle plot
(all species)
Component 1
Component 2

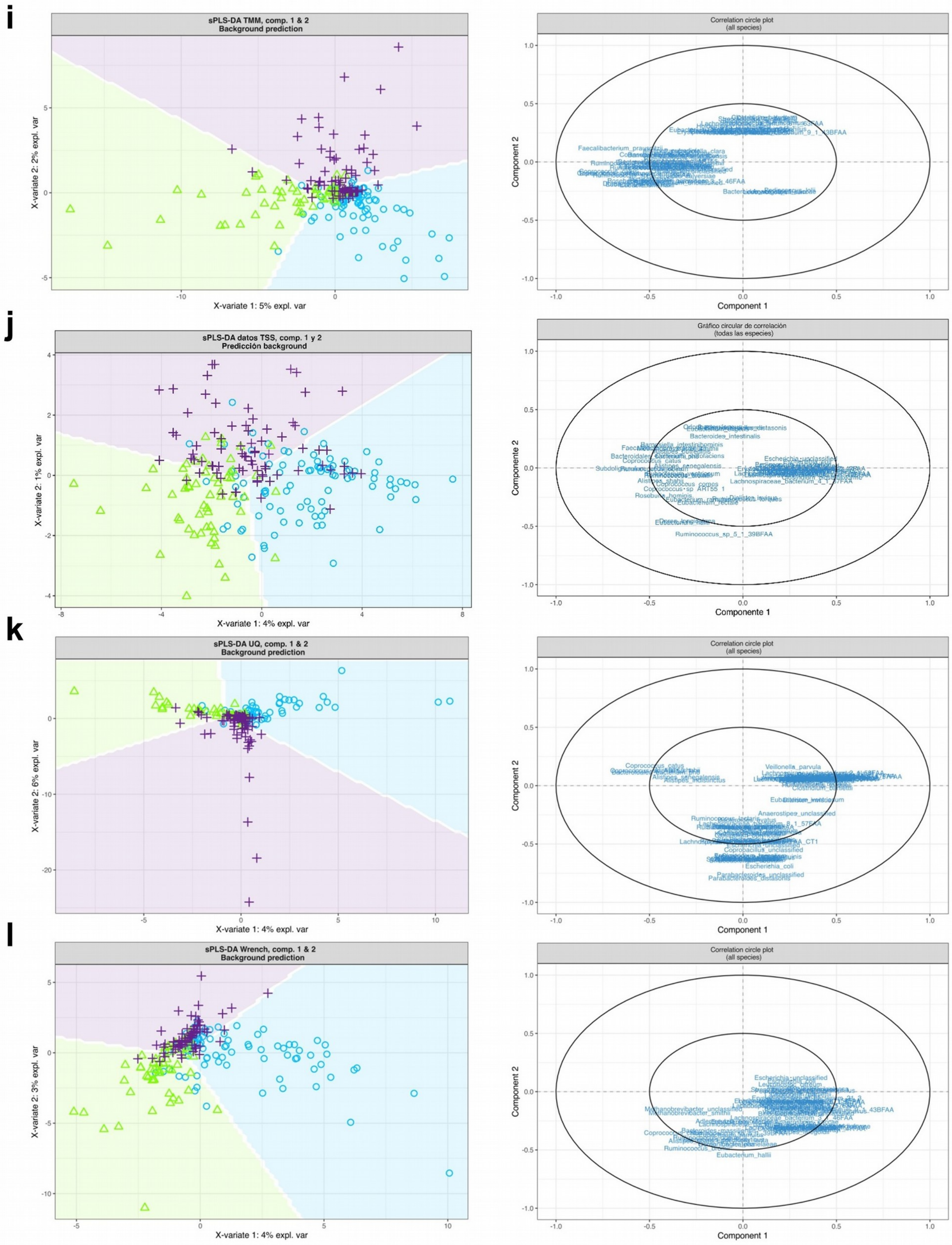
i
sPLS-DA TMM, comp. 1 & 2
Background prediction
X-variate 1: 5% expl. var
X-variate 2: 2% expl. var
Correlation circle plot
(all species)
Component 1
Component 2
Faecalibacterium_prausnitzii
j
sPLS-DA datos TSS, comp. 1 y 2
Predicción background
X-variate 1: 4% expl. var
X-variate 2: 1% expl. var
Gráfico circular de correlación
(todas las especies)
Componente 1
Componente 2
Bacteroides_intestinalis
Lachnospiraceae_bacterium_4_1_37FAA
Ruminococcus_sp_5_1_39BFAA
k
sPLS-DA UQ, comp. 1 & 2
Background prediction
X-variate 1: 4% expl. var
X-variate 2: 6% expl. var
Correlation circle plot
(all species)
Component 1
Component 2
Veillonella_parvula
Anaerostipes_unclassified
Coprobacillus_unclassified
Escherichia_coli
Parabacteroides_unclassified
l
sPLS-DA Wrench, comp. 1 & 2
Background prediction
X-variate 1: 4% expl. var
X-variate 2: 3% expl. var
Correlation circle plot
(all species)
Component 1
Component 2
Escherichia_unclassified
Eubacterium_hallii

**m**

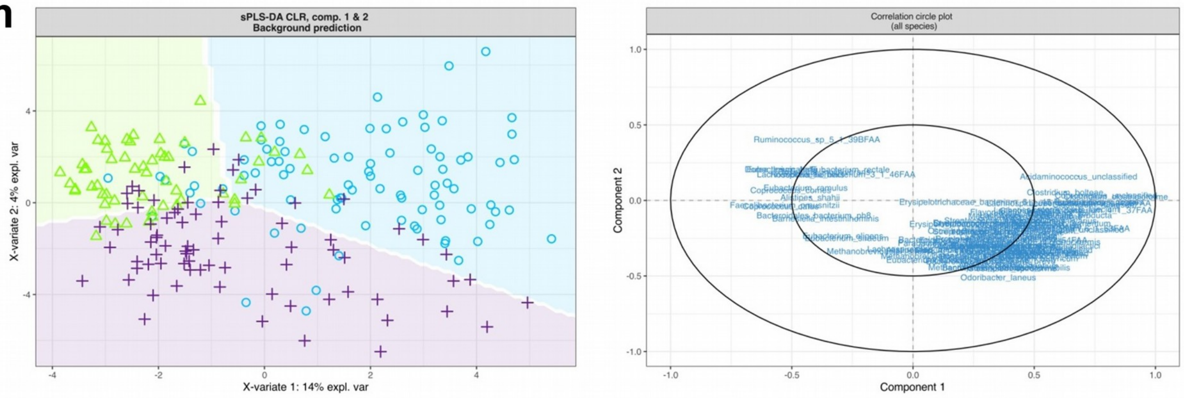

sPLS-DA CLR, comp. 1 & 2
Background prediction
X-variate 1: 14% expl. var
X-variate 2: 4% expl. var
Correlation circle plot
(all species)
Component 1
Component 2
Ruminococcus_sp_5_1_39BFAA
Acidaminococcus_unclassified
Clostridium_bolteae
Eubacterium_ramulus
Alistipes_shahii
Bacteroidales_bacterium_ph8
Eubacterium_eligens
Methanobrevibacter_smithii
Odoribacter_laneus
