## Supplementary Document 4 for "Alpha and beta-diversities performance comparison between different normalization methods and centered log-ratio transformation in a microbiome public dataset"

**Supplementary material 4**

Loading plots for each normalization methods and transformation method

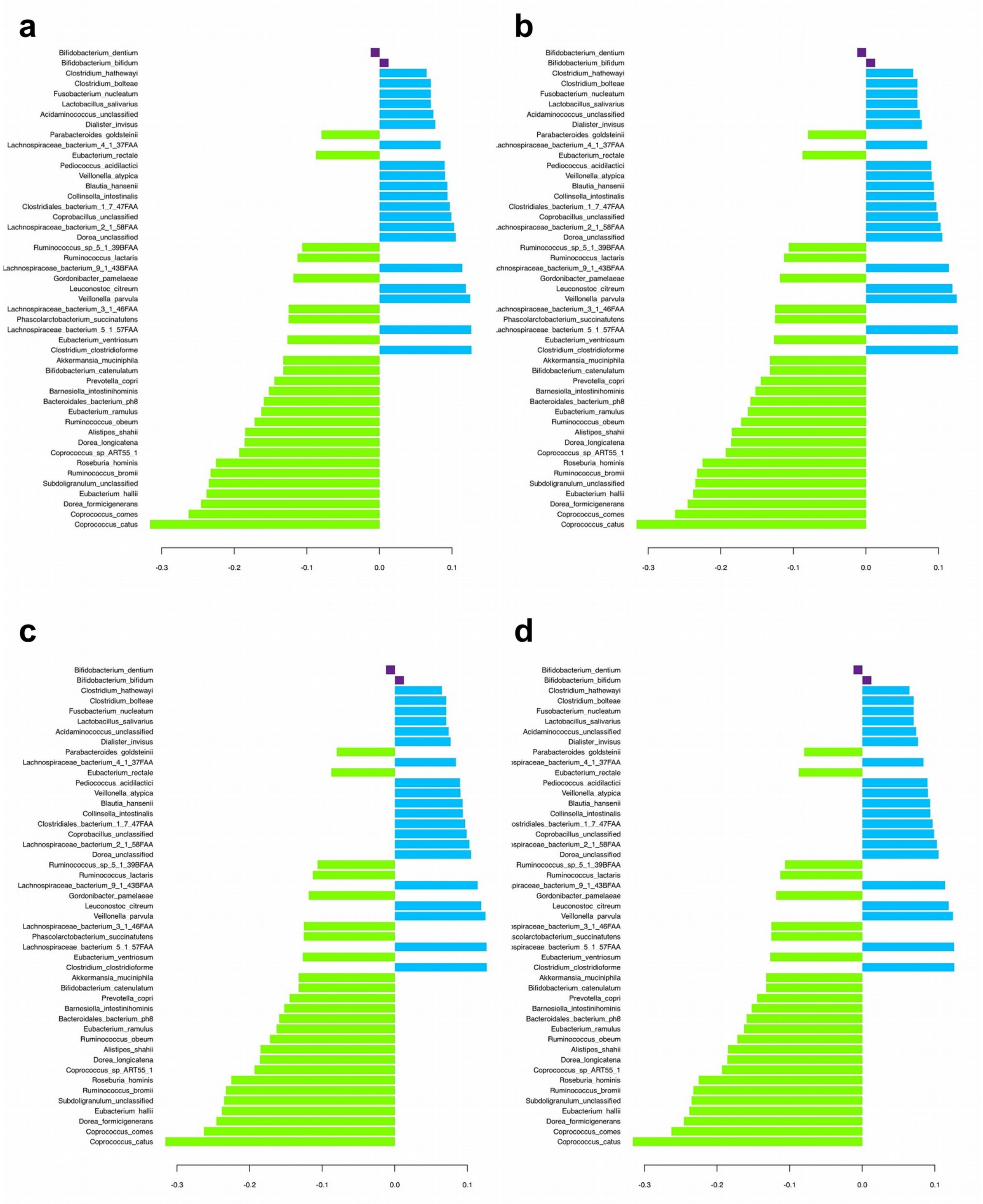

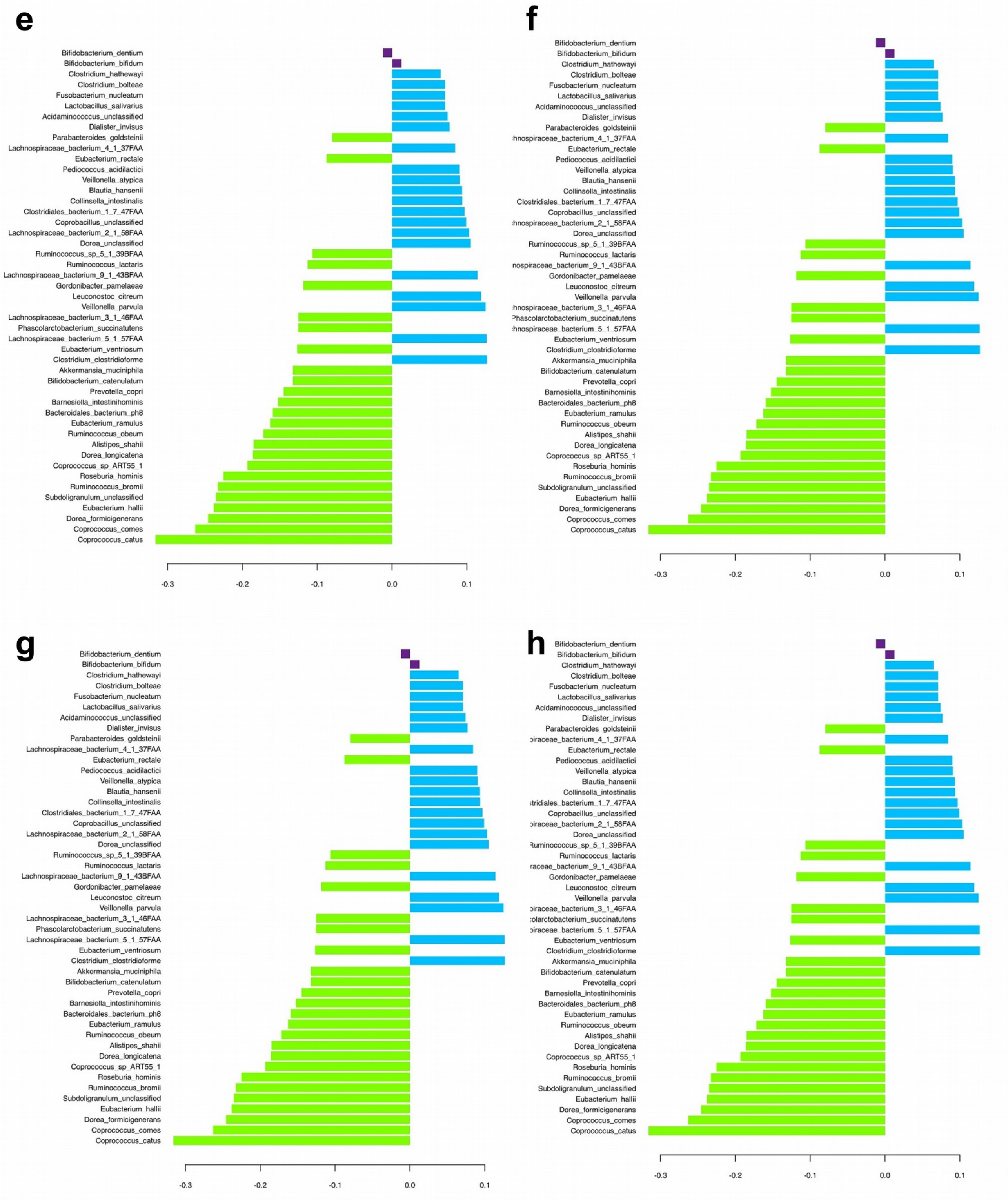

e
Bifidobacterium_dentium
Bifidobacterium_bifidum
Clostridium_hathewayi
Clostridium_bolteae
Fusobacterium_nucleatum
Lactobacillus_salivarius
Acidaminococcus_unclassified
Dialister_invisus
Parabacteroides_goldsteinii
Lachnospiraceae_bacterium_4_1_37FAA
Eubacterium_rectale
Pediococcus_acidilactici
Veillonella_atypica
Blautia_hansenii
Collinsella_intestinalis
Clostridiales_bacterium_1_7_47FAA
Coprobacillus_unclassified
Lachnospiraceae_bacterium_2_1_58FAA
Dorea_unclassified
Ruminococcus_sp_5_1_39BFAA
Ruminococcus_lactaris
Lachnospiraceae_bacterium_9_1_43BFAA
Gordonibacter_pamelaeae
Leuconostoc_citreum
Veillonella_parvula
Lachnospiraceae_bacterium_3_1_46FAA
Phascolarctobacterium_succinatutens
Lachnospiraceae_bacterium_5_1_57FAA
Eubacterium_ventriosum
Clostridium_clostridioforme
Akkermansia_muciniphila
Bifidobacterium_catenulatum
Prevotella_copri
Barnesiella_intestinihominis
Bacteroidales_bacterium_ph8
Eubacterium_ramulus
Ruminococcus_obeum
Alistipes_shahii
Dorea_longicatena
Coprococcus_sp_ART55_1
Roseburia_hominis
Ruminococcus_bromii
Subdoligranulum_unclassified
Eubacterium_hallii
Dorea_formicigenerans
Coprococcus_comes
Coprococcus_catus
-0.3
-0.2
-0.1
0.0
0.1
f
Bifidobacterium_dentium
Bifidobacterium_bifidum
Clostridium_hathewayi
Clostridium_bolteae
Fusobacterium_nucleatum
Lactobacillus_salivarius
Acidaminococcus_unclassified
Dialister_invisus
Parabacteroides_goldsteinii
hnospiraceae_bacterium_4_1_37FAA
Eubacterium_rectale
Pediococcus_acidilactici
Veillonella_atypica
Blautia_hansenii
Collinsella_intestinalis
Clostridiales_bacterium_1_7_47FAA
Coprobacillus_unclassified
hnospiraceae_bacterium_2_1_58FAA
Dorea_unclassified
Ruminococcus_sp_5_1_39BFAA
Ruminococcus_lactaris
nospiraceae_bacterium_9_1_43BFAA
Gordonibacter_pamelaeae
Leuconostoc_citreum
Veillonella_parvula
hnospiraceae_bacterium_3_1_46FAA
Phascolarctobacterium_succinatutens
hnospiraceae_bacterium_5_1_57FAA
Eubacterium_ventriosum
Clostridium_clostridioforme
Akkermansia_muciniphila
Bifidobacterium_catenulatum
Prevotella_copri
Barnesiella_intestinihominis
Bacteroidales_bacterium_ph8
Eubacterium_ramulus
Ruminococcus_obeum
Alistipes_shahii
Dorea_longicatena
Coprococcus_sp_ART55_1
Roseburia_hominis
Ruminococcus_bromii
Subdoligranulum_unclassified
Eubacterium_hallii
Dorea_formicigenerans
Coprococcus_comes
Coprococcus_catus
-0.3
-0.2
-0.1
0.0
0.1
g
Bifidobacterium_dentium
Bifidobacterium_bifidum
Clostridium_hathewayi
Clostridium_bolteae
Fusobacterium_nucleatum
Lactobacillus_salivarius
Acidaminococcus_unclassified
Dialister_invisus
Parabacteroides_goldsteinii
Lachnospiraceae_bacterium_4_1_37FAA
Eubacterium_rectale
Pediococcus_acidilactici
Veillonella_atypica
Blautia_hansenii
Collinsella_intestinalis
Clostridiales_bacterium_1_7_47FAA
Coprobacillus_unclassified
Lachnospiraceae_bacterium_2_1_58FAA
Dorea_unclassified
Ruminococcus_sp_5_1_39BFAA
Ruminococcus_lactaris
Lachnospiraceae_bacterium_9_1_43BFAA
Gordonibacter_pamelaeae
Leuconostoc_citreum
Veillonella_parvula
Lachnospiraceae_bacterium_3_1_46FAA
Phascolarctobacterium_succinatutens
Lachnospiraceae_bacterium_5_1_57FAA
Eubacterium_ventriosum
Clostridium_clostridioforme
Akkermansia_muciniphila
Bifidobacterium_catenulatum
Prevotella_copri
Barnesiella_intestinihominis
Bacteroidales_bacterium_ph8
Eubacterium_ramulus
Ruminococcus_obeum
Alistipes_shahii
Dorea_longicatena
Coprococcus_sp_ART55_1
Roseburia_hominis
Ruminococcus_bromii
Subdoligranulum_unclassified
Eubacterium_hallii
Dorea_formicigenerans
Coprococcus_comes
Coprococcus_catus
-0.3
-0.2
-0.1
0.0
0.1
h
Bifidobacterium_dentium
Bifidobacterium_bifidum
Clostridium_hathewayi
Clostridium_bolteae
Fusobacterium_nucleatum
Lactobacillus_salivarius
Acidaminococcus_unclassified
Dialister_invisus
Parabacteroides_goldsteinii
piraceae_bacterium_4_1_37FAA
Eubacterium_rectale
Pediococcus_acidilactici
Veillonella_atypica
Blautia_hansenii
Collinsella_intestinalis
stridiales_bacterium_1_7_47FAA
Coprobacillus_unclassified
piraceae_bacterium_2_1_58FAA
Dorea_unclassified
Ruminococcus_sp_5_1_39BFAA
Ruminococcus_lactaris
raceae_bacterium_9_1_43BFAA
Gordonibacter_pamelaeae
Leuconostoc_citreum
Veillonella_parvula
piraceae_bacterium_3_1_46FAA
colarctobacterium_succinatutens
piraceae_bacterium_5_1_57FAA
Eubacterium_ventriosum
Clostridium_clostridioforme
Akkermansia_muciniphila
Bifidobacterium_catenulatum
Prevotella_copri
Barnesiella_intestinihominis
Bacteroidales_bacterium_ph8
Eubacterium_ramulus
Ruminococcus_obeum
Alistipes_shahii
Dorea_longicatena
Coprococcus_sp_ART55_1
Roseburia_hominis
Ruminococcus_bromii
Subdoligranulum_unclassified
Eubacterium_hallii
Dorea_formicigenerans
Coprococcus_comes
Coprococcus_catus
-0.3
-0.2
-0.1
0.0
0.1

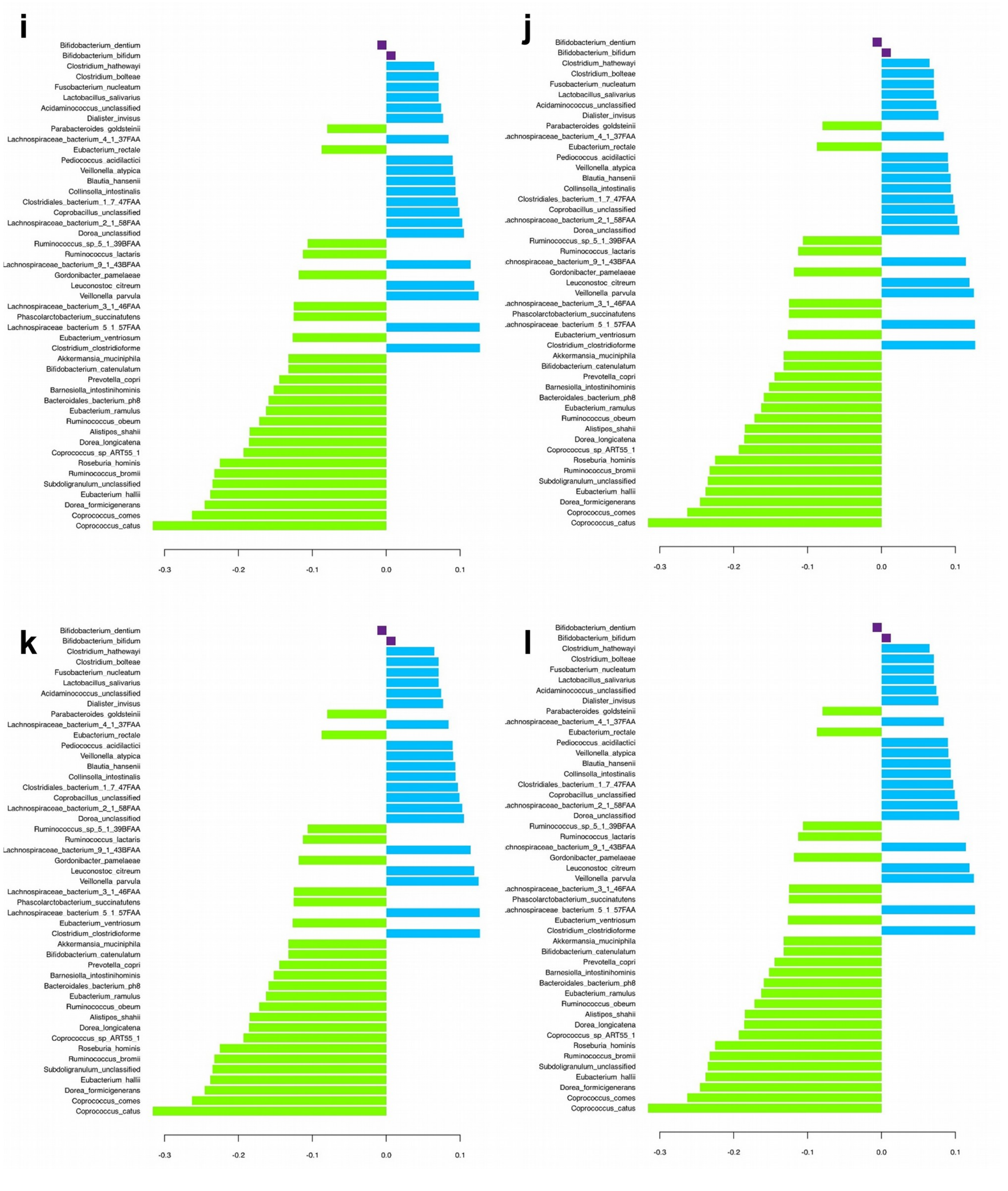
i
j
k
l
Bifidobacterium_dentium
Bifidobacterium_bifidum
Clostridium_hathewayi
Clostridium_bolteae
Fusobacterium_nucleatum
Lactobacillus_salivarius
Acidaminococcus_unclassified
Dialister_invisus
Parabacteroides_goldsteinii
Lachnospiraceae_bacterium_4_1_37FAA
Eubacterium_rectale
Pediococcus_acidilactici
Veillonella_atypica
Blautia_hansenii
Collinsella_intestinalis
Clostridiales_bacterium_1_7_47FAA
Coprobacillus_unclassified
Lachnospiraceae_bacterium_2_1_58FAA
Dorea_unclassified
Ruminococcus_sp_5_1_39BFAA
Ruminococcus_lactaris
Lachnospiraceae_bacterium_9_1_43BFAA
Gordonibacter_pamelaeae
Leuconostoc_citreum
Veillonella_parvula
Lachnospiraceae_bacterium_3_1_46FAA
Phascolarctobacterium_succinatutens
Lachnospiraceae_bacterium_5_1_57FAA
Eubacterium_ventriosum
Clostridium_clostridioforme
Akkermansia_muciniphila
Bifidobacterium_catenulatum
Prevotella_copri
Barnesiella_intestinihominis
Bacteroidales_bacterium_ph8
Eubacterium_ramulus
Ruminococcus_obeum
Alistipes_shahii
Dorea_longicatena
Coprococcus_sp_ART55_1
Roseburia_hominis
Ruminococcus_bromii
Subdoligranulum_unclassified
Eubacterium_hallii
Dorea_formicigenerans
Coprococcus_comes
Coprococcus_catus
-0.3
-0.2
-0.1
0.0
0.1

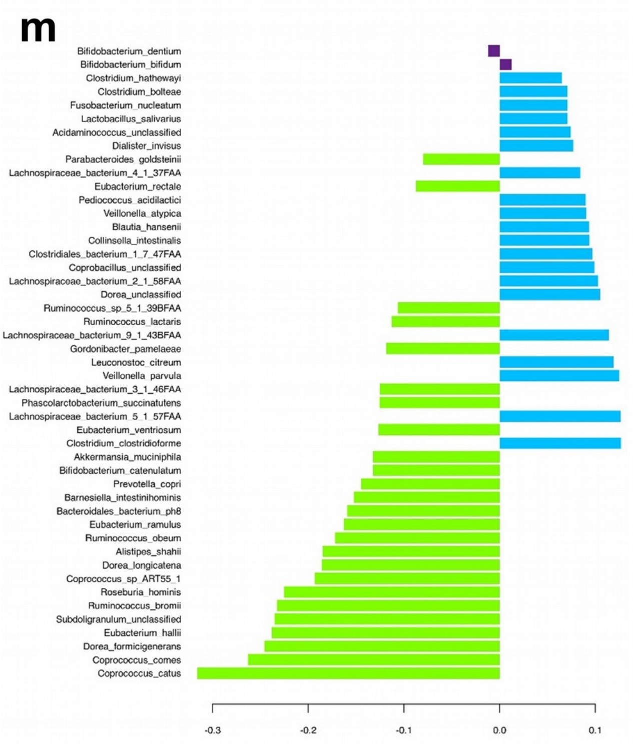
m
Bifidobacterium_dentium
Bifidobacterium_bifidum
Clostridium_hathewayi
Clostridium_bolteae
Fusobacterium_nucleatum
Lactobacillus_salivarius
Acidaminococcus_unclassified
Dialister_invisus
Parabacteroides_goldsteinii
Lachnospiraceae_bacterium_4_1_37FAA
Eubacterium_rectale
Pediococcus_acidilactici
Veillonella_atypica
Blautia_hansenii
Collinsella_intestinalis
Clostridiales_bacterium_1_7_47FAA
Coprobacillus_unclassified
Lachnospiraceae_bacterium_2_1_58FAA
Dorea_unclassified
Ruminococcus_sp_5_1_39BFAA
Ruminococcus_lactaris
Lachnospiraceae_bacterium_9_1_43BFAA
Gordonibacter_pamelaeae
Leuconostoc_citreum
Veillonella_parvula
Lachnospiraceae_bacterium_3_1_46FAA
Phascolarctobacterium_succinatutens
Lachnospiraceae_bacterium_5_1_57FAA
Eubacterium_ventriosum
Clostridium_clostridioforme
Akkermansia_muciniphila
Bifidobacterium_catenulatum
Prevotella_copri
Barnesiella_intestinihominis
Bacteroidales_bacterium_ph8
Eubacterium_ramulus
Ruminococcus_obeum
Alistipes_shahii
Dorea_longicatena
Coprococcus_sp_ART55_1
Roseburia_hominis
Ruminococcus_bromii
Subdoligranulum_unclassified
Eubacterium_hallii
Dorea_formicigenerans
Coprococcus_comes
Coprococcus_catus
-0.3
-0.2
-0.1
0.0
0.1
